# An integrative systems-biology approach defines mechanisms of Alzheimer’s disease neurodegeneration

**DOI:** 10.1101/2024.03.17.585262

**Authors:** Matthew J Leventhal, Camila A Zanella, Byunguk Kang, Jiajie Peng, David Gritsch, Zhixiang Liao, Hassan Bukhari, Tao Wang, Ping-Chieh Pao, Serwah Danquah, Joseph Benetatos, Ralda Nehme, Samouil Farhi, Li-Huei Tsai, Xianjun Dong, Clemens R Scherzer, Mel B Feany, Ernest Fraenkel

**Affiliations:** MIT Ph.D. Program in Computational and Systems Biology, Cambridge, MA, USA; Department of Biological Engineering, Massachusetts Institute of Technology, Cambridge, MA, USA; Broad Institute of Harvard and MIT, Cambridge, MA, USA; Department of Pathology, Brigham and Women’s Hospital and Harvard Medical School, Boston, MA, USA; Spatial Technology Platform, Broad Institute of Harvard and MIT, Cambridge, MA USA; Precision Neurology Program, Brigham and Women’s Hospital and Harvard Medical school, Boston, MA, USA; APDA Center for Advanced Parkinson’s Disease Research, Brigham and Women’s Hospital and Harvard Medical School, Boston, MA, USA; Picower Institute for Learning and Memory, Massachusetts Institute of Technology, Cambridge, Massachusetts, USA; Department of Brain and Cognitive Sciences, Massachusetts Institute of Technology, Cambridge, Massachusetts, USA; Stanley Center for Psychiatric Research, Broad Institute of Harvard and MIT, Cambridge, MA, USA; Present address: School of Computer Science, Northwestern Polytechnical University, Xi’an, China; Present address: Stephen and Denise Adams Center of Yale School of Medicine, CT, USA

## Abstract

Despite years of intense investigation, the mechanisms underlying neuronal death in Alzheimer’s disease, the most common neurodegenerative disorder, remain incompletely understood. To define relevant pathways, we integrated the results of an unbiased, genome-scale forward genetic screen for age-associated neurodegeneration in *Drosophila* with human and *Drosophila* Alzheimer’s disease-associated multi-omics. We measured proteomics, phosphoproteomics, and metabolomics in *Drosophila* models of Alzheimer’s disease and identified Alzheimer’s disease human genetic variants that modify expression in disease-vulnerable neurons. We used a network optimization approach to integrate these data with previously published Alzheimer’s disease multi-omic data. We computationally predicted and experimentally demonstrated how *HNRNPA2B1* and *MEPCE* enhance tau-mediated neurotoxicity. Furthermore, we demonstrated that the screen hits *CSNK2A1* and *NOTCH1* regulate DNA damage in *Drosophila* and human iPSC-derived neural progenitor cells. Our work identifies candidate pathways that could be targeted to ameliorate neurodegeneration in Alzheimer’s disease.

## Introduction

Neurodegenerative diseases are characterized by a progressive loss of neurons and preferentially affect older individuals. As the global population ages, there is an increasing imperative to understand and design effective therapies for neurodegenerative disorders. Brains from patients with Alzheimer’s disease, the most common neurodegenerative disorder, show pathological aggregation and deposition of extracellular amyloid β plaques and intracellular neurofibrillary tangles comprised of tau protein^1–3^. Amyloid β plaques are predominantly made up of 42-amino acid amyloid β peptides (amyloid β_1-42_), which are processed from the larger amyloid precursor protein (APP)^4^ through the action of the gamma-secretase complex. The presenilin proteins comprise the protease subunit of gamma-secretase. Mutations in *APP* and in the genes encoding the presenilins give rise to fully penetrant, though rare, familial Alzheimer’s disease. Mutations in *MAPT*, the gene encoding the microtubule-associated protein tau, have not been discovered in Alzheimer’s disease patients. However, wild-type tau is the major aggregating protein in a group of sporadic neurodegenerative disorders characterized by tau deposition in neurofibrillary aggregates, termed tauopathies. Further, missense mutations in *MAPT* cause fully penetrant, severe forms of familial neurodegeneration, strongly linking tau to aging-dependent neurodegeneration^5^.

Discovery of single gene mutations leading to familial forms of neurodegenerative disease provided a critical starting point for developing animal models and understanding underlying pathophysiology. More recently, data derived from high content approaches has added to clues from classical genetics. Genome-wide association studies (GWAS), transcriptomic analysis, and quantitative trait locus (QTL) analysis have identified genetic risk factors and associated molecular changes underlying Alzheimer’s disease in the brain at bulk and single-neuron resolution^6–13^.

Despite the wealth of human pathological and genetic data, the pathways through which both single gene mutations and QTL-associated molecular changes impact neurodegenerative disease pathogenesis remain incompletely defined. Complementary approaches are thus needed to define the full set of mechanisms mediating neurodegenerative disease pathogenesis. A more complete understanding of cell death pathways should provide an important new set of therapeutic targets in Alzheimer’s disease and related age-dependent neurodegenerative disorders^14^.

Forward screens in organisms, like *Drosophila*, with relatively short lifespan, well-developed experimental tools, and conserved neuronal biology, represent a potentially valuable method for discovering new pathways mediating neurodegeneration. Indeed, prior genetic screens in tauopathy model flies have identified a number of pathways mediating toxicity of pathological human tau that are conserved in vertebrate systems^15–28^. Similarly, genetic screens in otherwise wild type flies have identified mutants causing progressive neurodegeneration with relevance to human disease, but these efforts have been relatively modest in scale^29–33^. Here we have employed unbiased forward genetic screening at genome-scale in *Drosophila* to define mechanisms required for maintenance of aging adult neurons.

We then used a multi-omic integration approach to relate the hits from our model organism screen to human disease and identify the pathways that influence age-associated neurodegeneration. We measured proteomics, phosphoproteomics and metabolomics in transgenic *Drosophila* models of human amyloid β and tau to identify molecular changes associated with Alzheimer’s disease toxic proteins (Fig. 1). To determine how our transgenic *Drosophila* RNAi screen and the other model organism data were related to human Alzheimer’s disease patients, we generated RNA-sequencing (RNA-seq) data from pyramidal neuron-enriched populations from the temporal cortex using laser-capture microdissection^34–38^ (Fig. 1). We identified fine-mapped expression QTLs (eQTLs) and the eQTL-associated genes (eGenes) in neurons vulnerable to disease pathology to find patterns of gene expression associated with human genetic risk factors of Alzheimer’s disease. Next, we integrated these multi-species, multi-omic data with a previously published genome-scale screen for tau-mediated neurotoxicity^15^, existing human Alzheimer’s disease GWAS hits, proteomics, and metabolomics ^7,9,15,39,40^ using advanced network modeling approaches (Fig. 1)^41,42^.

**Fig. 1:**
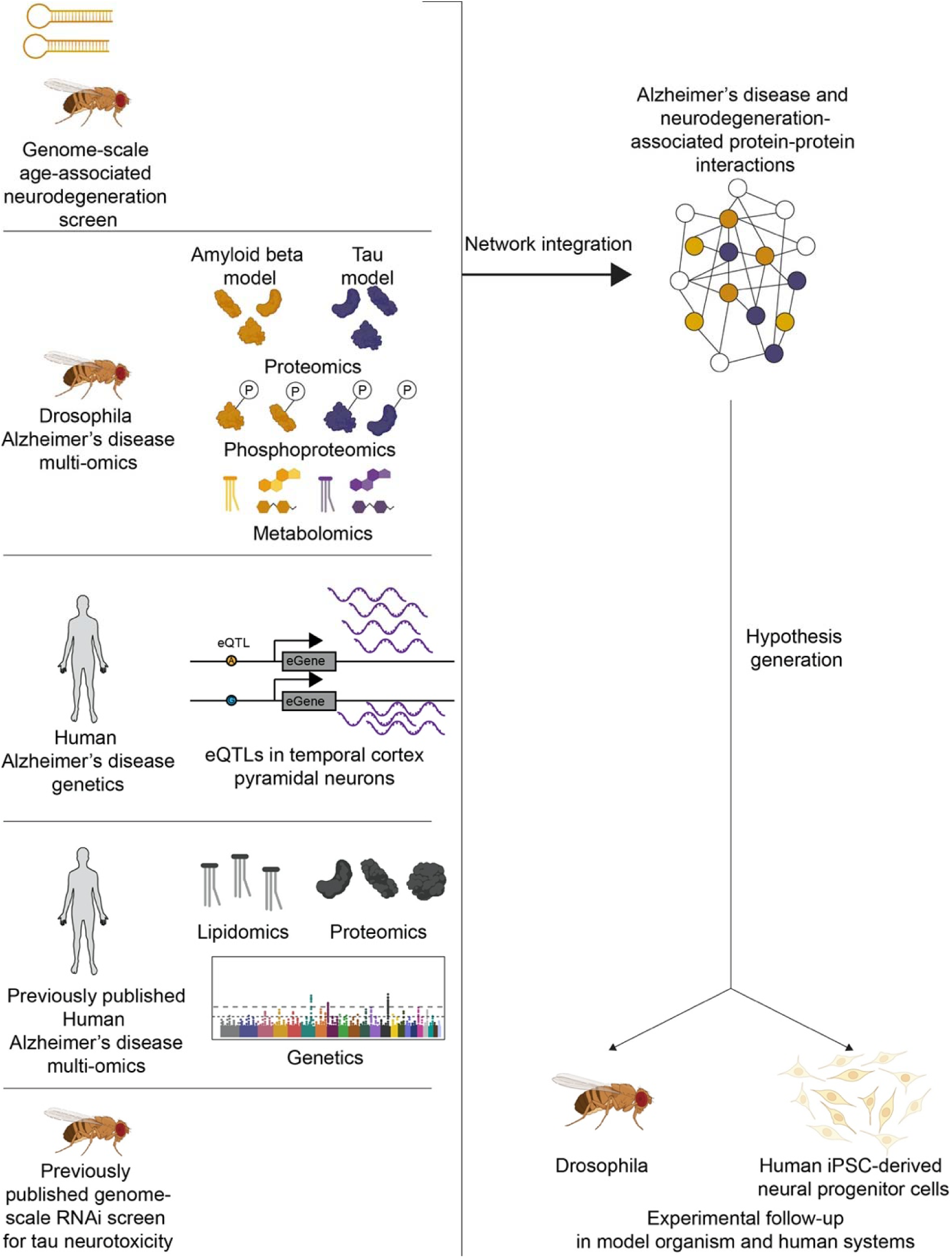
Overview of analytical framework for multi-omic integration to study the biological processes underlying neurodegeneration. We performed a forward genetic screen for age-associated neurodegeneration in *Drosophila*. We measured proteomics, phosphoproteomics and metabolomics in amyloid β (gold) and tau (purple) models of Alzheimer’s disease and performed an eQTL meta-analysis of previous Alzheimer’s disease studies. We used a network integration model to integrate these new data with previously published human proteomics, human genetics, human lipidomics, and *Drosophila* modifiers of tau-mediated neurotoxicity. We tested hypotheses generated from this network model in *Drosophila* and human iPSC-derived neural progenitor cells. Icons created with Biorender.com.

Based on our integrated model, we nominated genes and pathways that contribute to age-associated neurodegeneration in Alzheimer’s disease. We experimentally tested the predicted functional effects of knockdown of proposed targets in flies and in human induced pluripotent stem cells. Specifically, we demonstrate that the human Alzheimer’s disease genetic risk factor *MEPCE* and neurodegeneration screen hit *HNRNPA2B1* regulate tau-mediated neurotoxicity. Furthermore, we show in flies and iPSC-derived neural progenitor cells that *NOTCH1* and *CSNK2A1* regulate the DNA damage response, suggesting pathways through which these genes enhance neurodegeneration.

## Results

### A genome-scale, forward genetic screen for neurodegeneration in Drosophila

We performed a genome-scale, forward genetic screen in *Drosophila* to identify genes required for maintenance of viability of aging neurons *in vivo*. We used transgenic RNAi to knock down 5,261 fly genes in neurons using the UAS-GAL4 bipartite expression system and the widely used *elav-GAL4* driver, which mediates expression in the pattern of the pan-neuronal gene *elav*^43,44^. Genes were knocked down based on availability of transgenic RNAi lines from the public Bloomington *Drosophila* Stock Center and were not otherwise preselected. Adult flies with neuronal gene knockdown were aged for 30 days and brain integrity was assessed on tissue sections representing the entire brain. Scoring was performed in a blinded fashion. Genes were identified as hits in the screen if there was neuronal loss or vacuolation in the context of a properly developed brain. Neurodegeneration is frequently accompanied by vacuolation of the brain in flies and is commonly viewed as a sensitive and specific measure of neurodegeneration in the brain aging and neurodegenerative disease models ^20,29,33,45–57^. Vacuoles often occur in a range of human neurodegenerative disease as well, including Alzheimer’s disease and related tauopathies^58–64^. We identified 198 genes that promoted age-associated neurodegeneration in *Drosophila* after knockdown (Table 1). Strikingly, we recovered orthologs of both *APP* (*Appl*) and the presenilins (*Psn*), genes mutated in the two monogenic causes of familial Alzheimer’s disease. Our findings are consistent with demonstration of age-dependent neurodegeneration in *Appl* and *Psn* mutants in other studies^28,65^. In addition to *Drosophila APP* and *Psn* we also recovered fly orthologs of genes mutated in monogenic forms of Parkinson’s disease (*PLA2G6/iPLA2-VIA*), amyotrophic lateral sclerosis (*SOD1/Sod1, VAPB/Vap33*), hereditary spastic paraparesis (*VPS37A/Vps37A*), and mitochondrial encephalomyopathy (*COX10/Cox10, NDUFS7/ND-20*, *NDUFV1/ND-51, MSTO1/Mst, TTC19/Ttc19*). Further connecting the results of our screen with neurodegeneration we observed an enrichment for ubiquitin-associated pathways (Benjamini-Hochberg FDR-adjusted p-value=1.48*10^-3^).

**Table 1.**
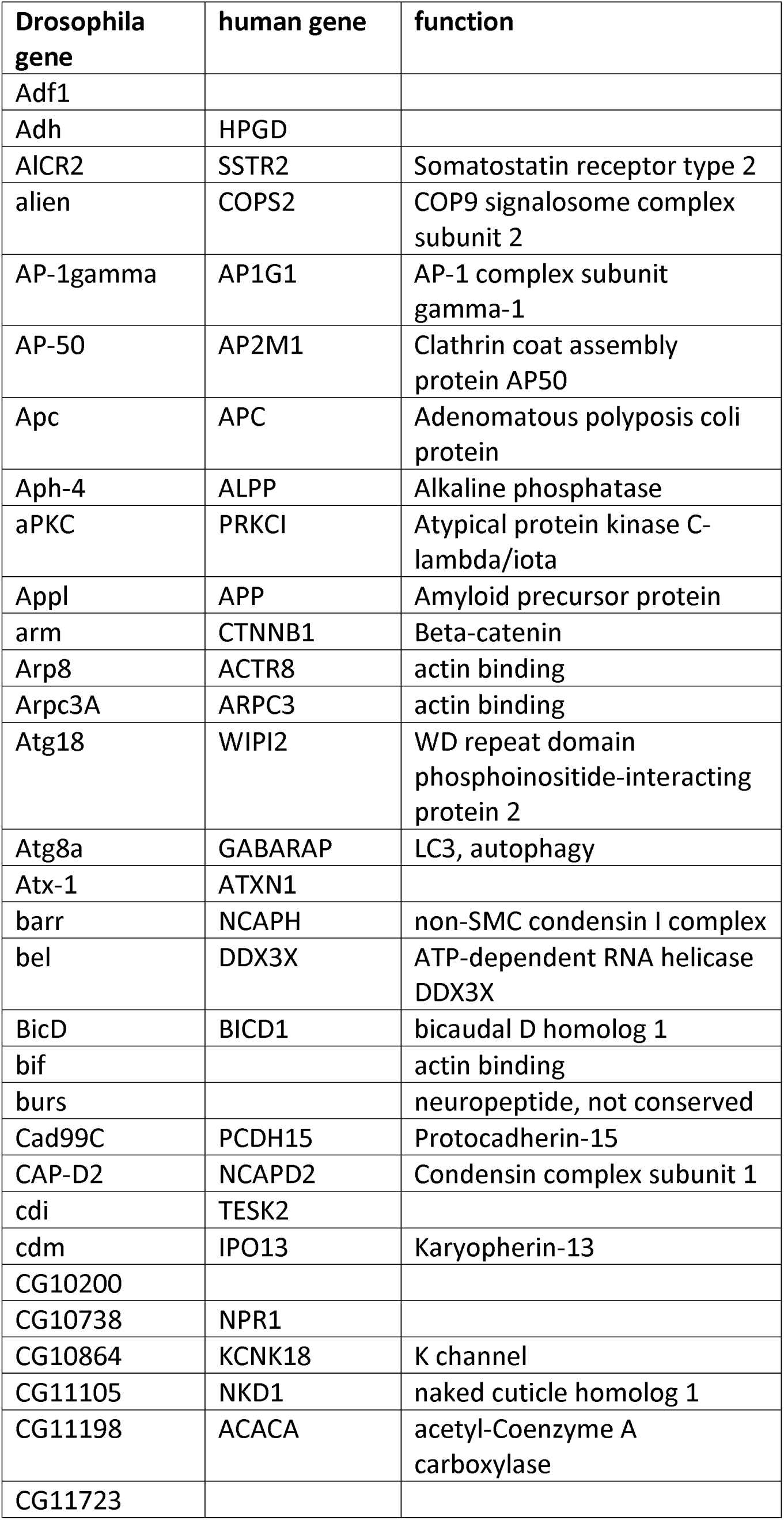
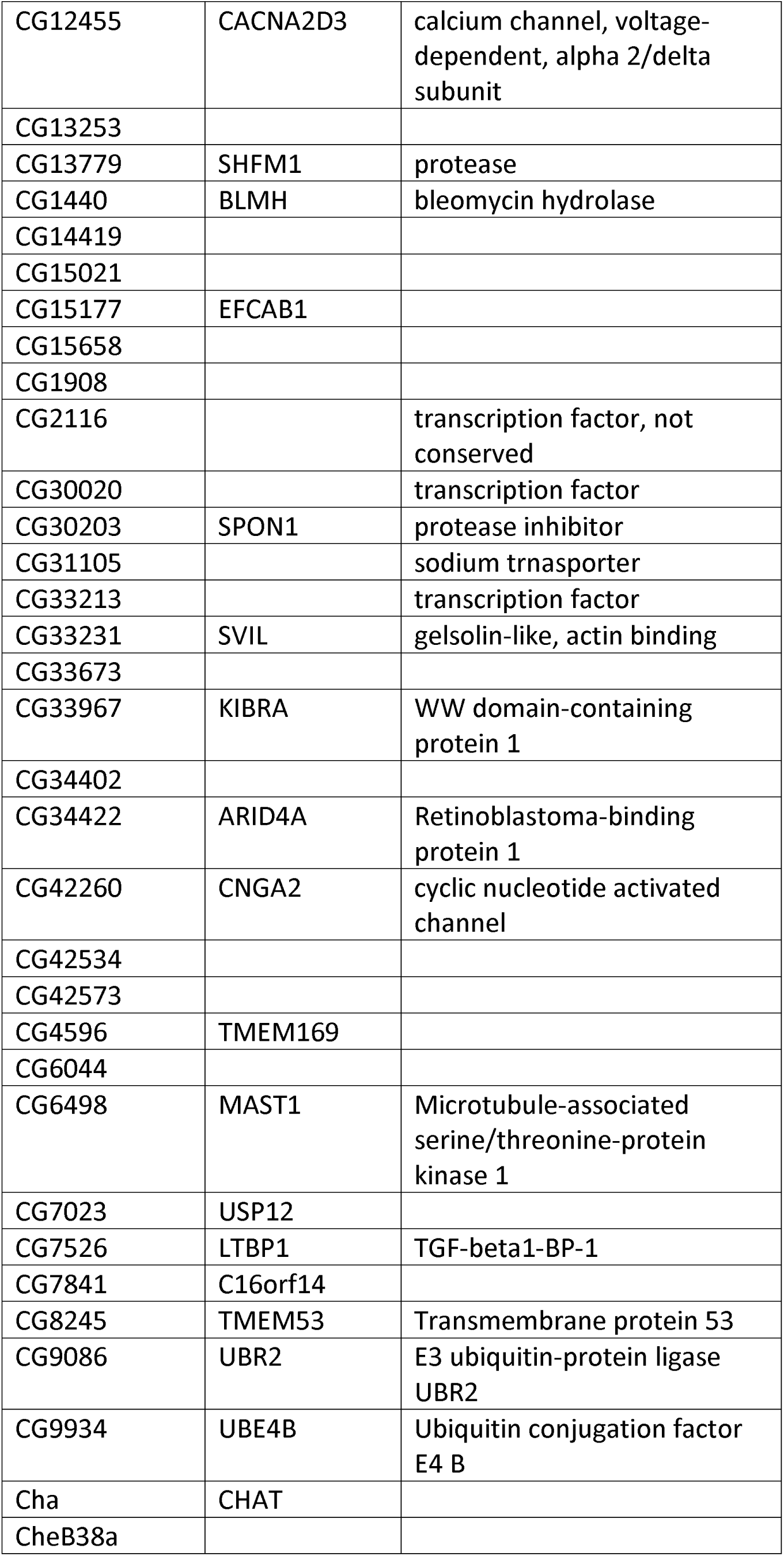
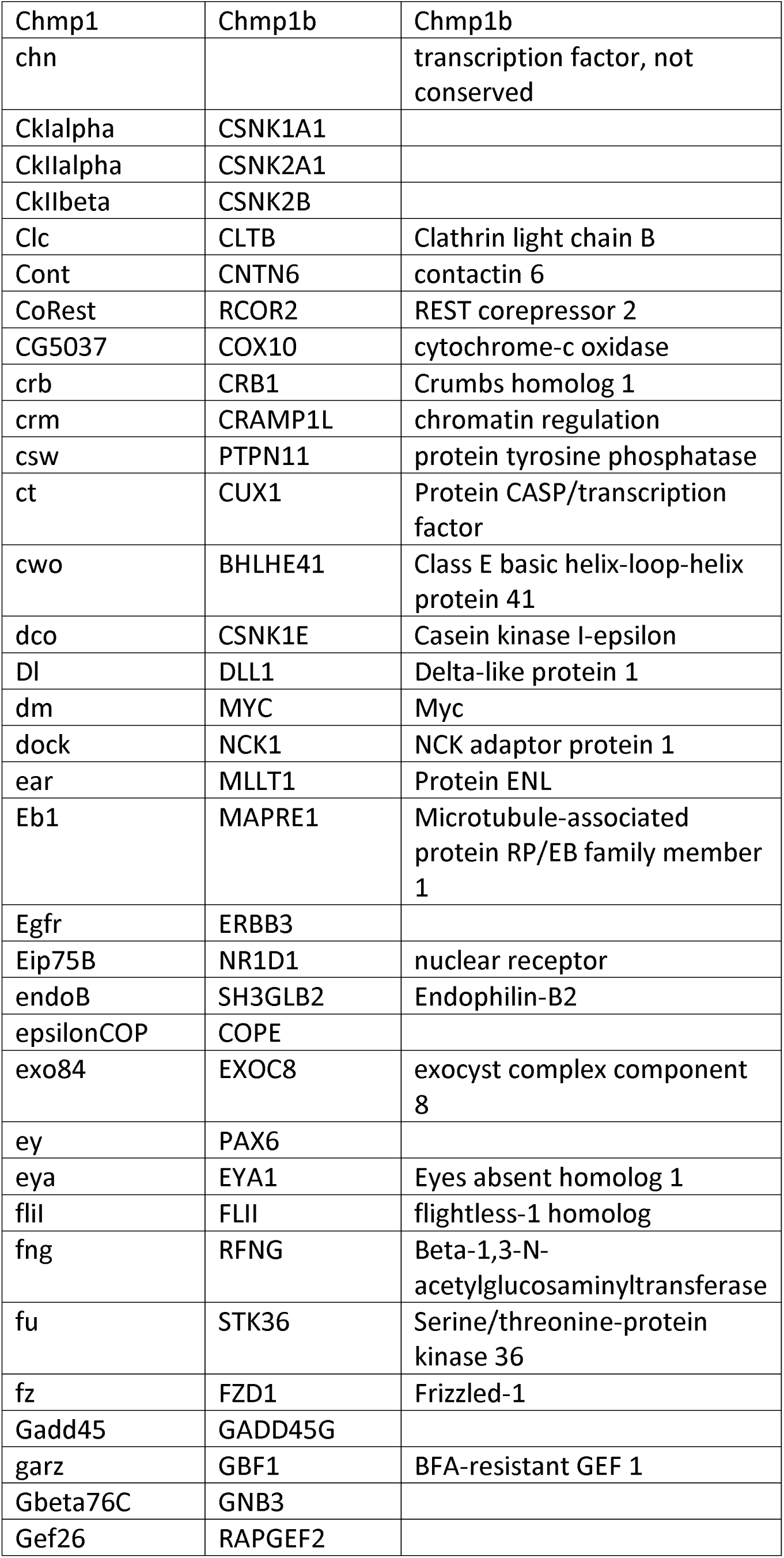
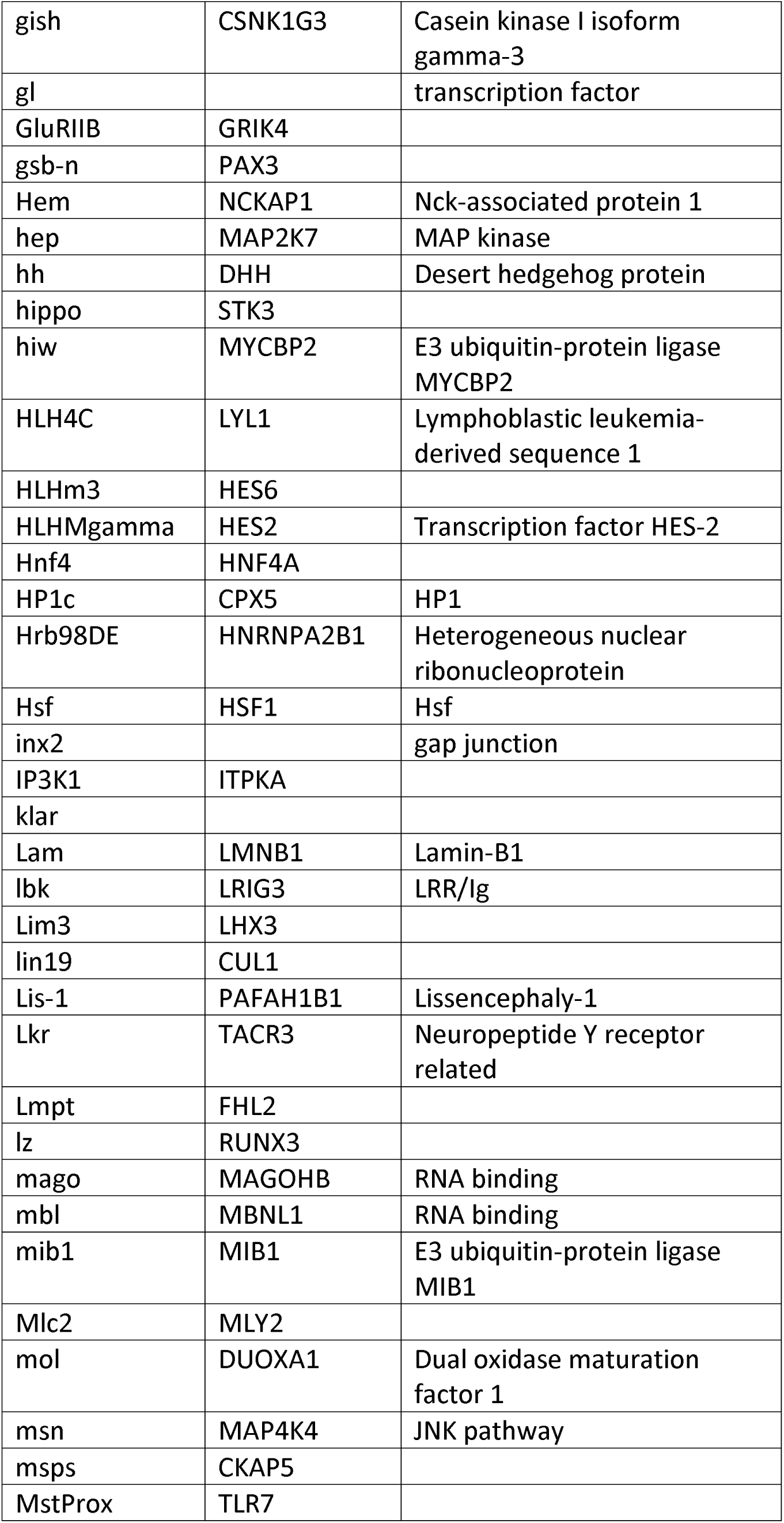
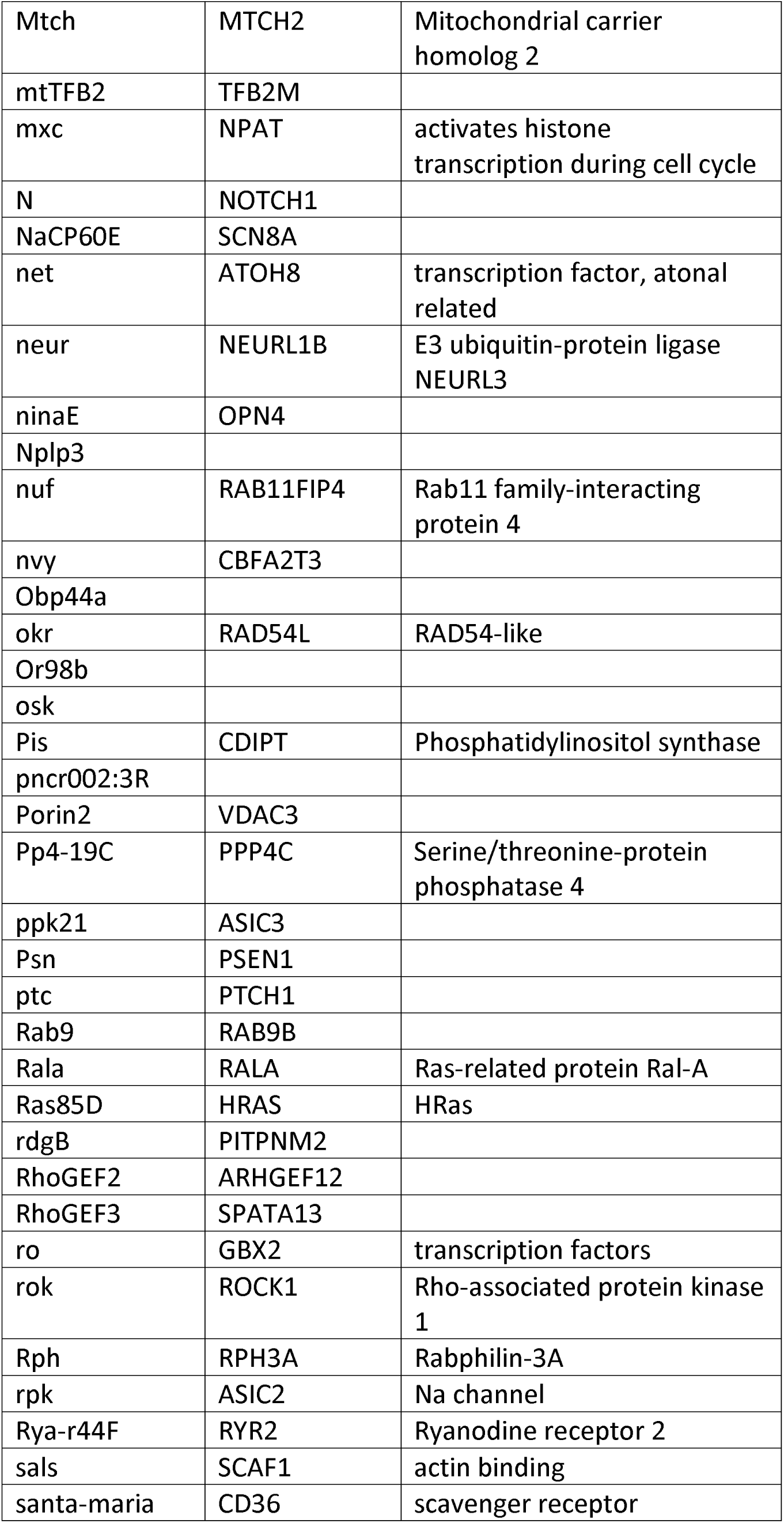
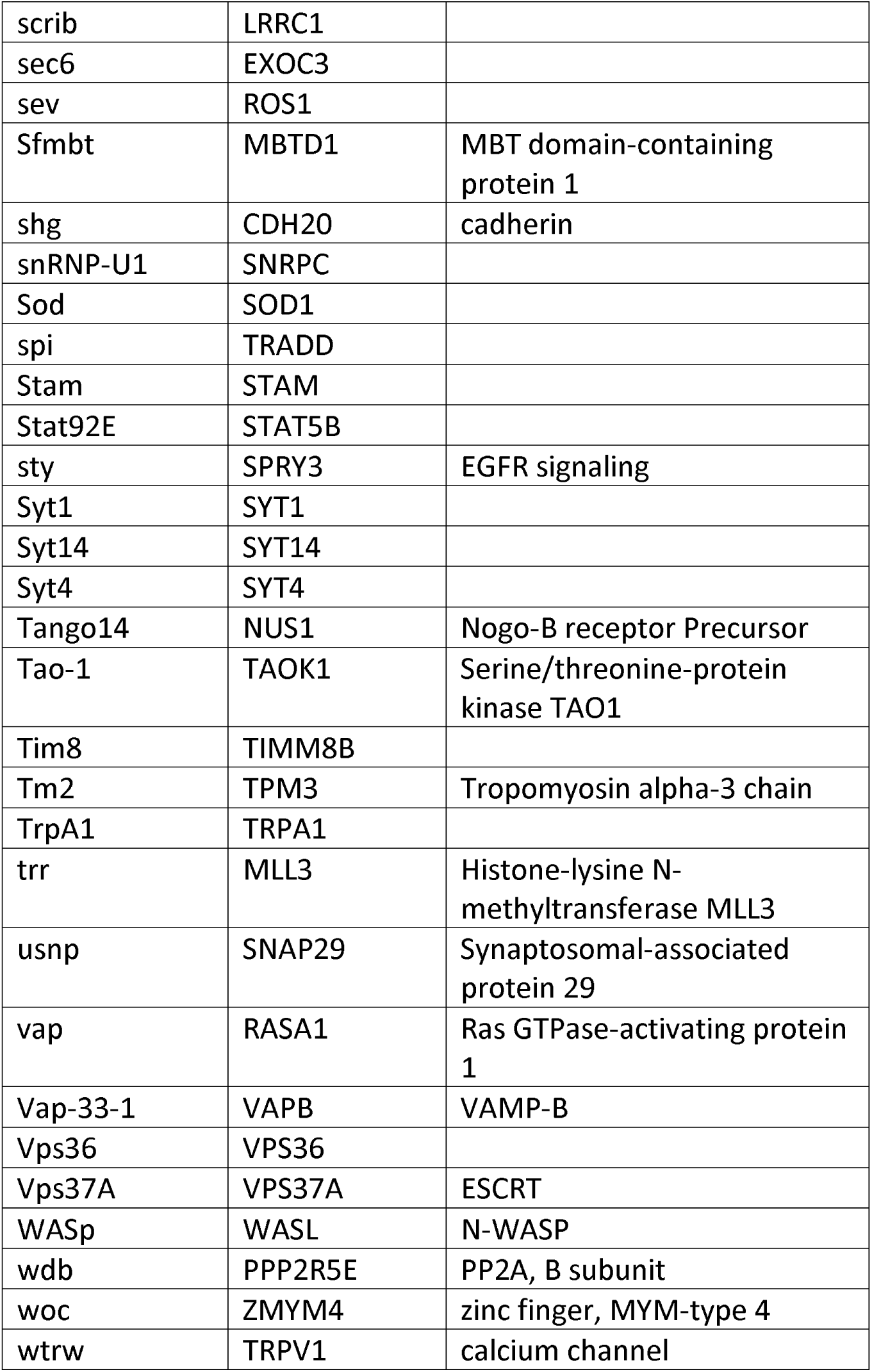
Hits from the age-associated neurodegeneration screen. : List of *Drosophila* genes and human orthologs that were hits from the screen for age-associated neurodegeneration. Human orthologs of genetic screen hits in *Drosophila* were inferred using DIOPT^126^.

### Gene expression of the human orthologs of the neurodegeneration screen hits declines with age and Alzheimer’s disease in the human brain

To assess more broadly if *Drosophila* screen hits were associated with human neurodegeneration we analyzed RNA-seq data from 2642 human post-mortem brain tissues from the Genotype-Tissue Expression (GTEx) project. We assessed whether there was a significant association between the mean expression of the human orthologs of the neurodegeneration screen hits and age in these human brains. We found that the average expression of neurodegeneration screen hits was negatively associated with chronological age (Fig. 2a, p=1.14*10^-5^). There was a stronger negative association between average gene expression and age for the neurodegeneration screen hits than the average expression of all protein-coding genes (Fig. 2a, all protein-coding genes: R=0.12, neurodegeneration screen hits: R=0.15). We found there was a significant difference in the slopes of the regression lines showing the relationship between gene expression and age for neurodegeneration screen hits compared to that of all protein-coding genes (p=7.38*10^-6^). We subsequently ranked all genes by the regression coefficients measuring the relationship between gene expression and age. We performed Gene Set Enrichment Analysis on this ranked list to identify which pathways had significant changes in gene expression with respect to age. Our analysis showed a negative association between the expression of screen hits and age (Fig. 2b, Benjamini-Hochberg FDR-adjusted p-value<0.1). We note that the average expression of neurodegeneration screen hits is significantly greater than that of all protein-coding genes (Wilcoxon rank-sum test, p<1*10^-16^). To assess the robustness of our results, we performed permutation tests by randomly shuffling the patient ages. Not a single permutation out of 10,000 iterations had a more significant association between age and gene expression of the screen hits, suggesting that this result is specific to chronological age in humans.

**Fig. 2:**
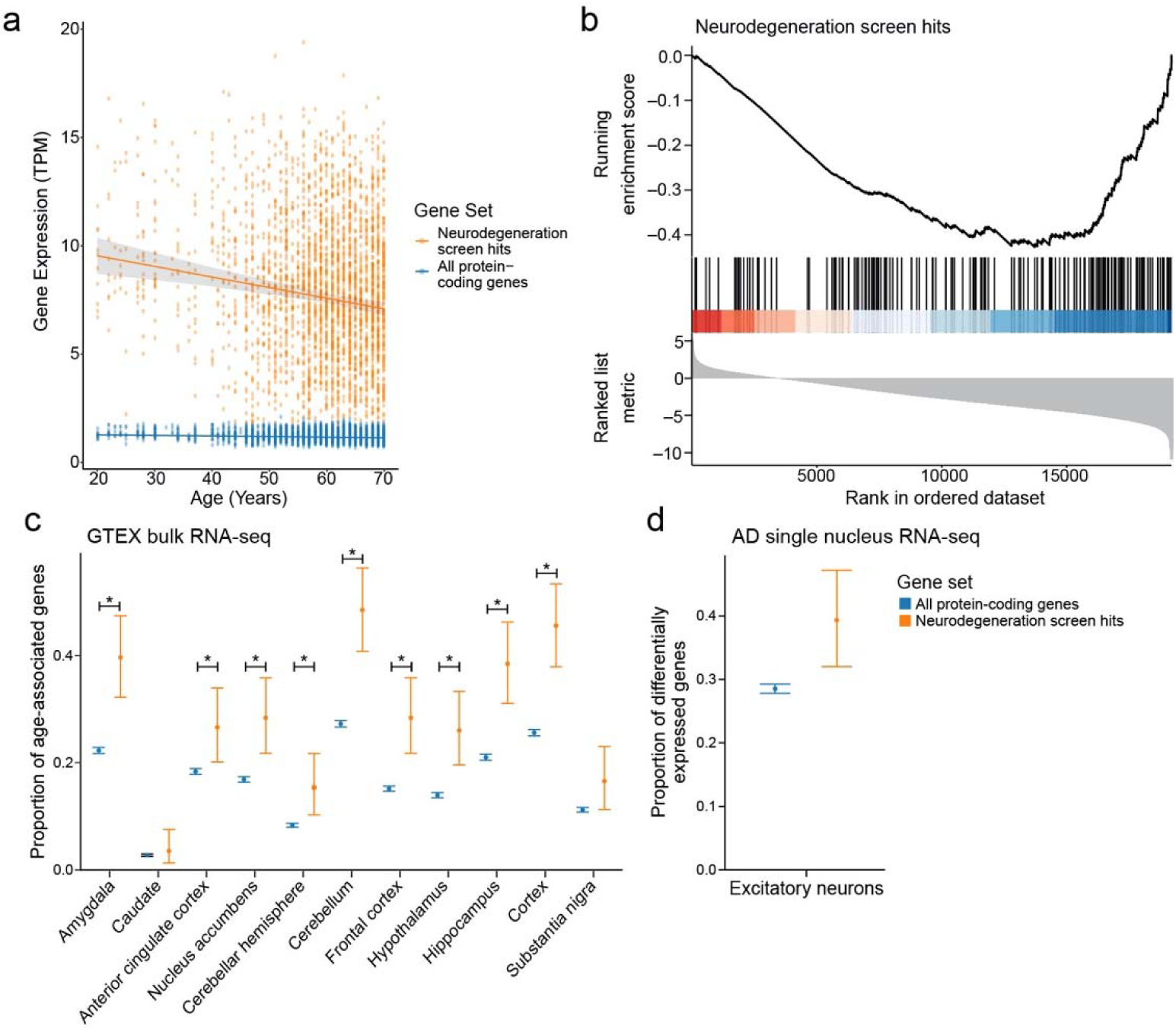
a) Geometric mean expression in transcripts per million (TPM) of neurodegeneration screen hits (neurodegeneration genes, orange) and all protein-coding genes in the Genotype-Tissue Expression (GTEx) shows that the expression of neurodegeneration screen hits declines with age in human brain tissues (all protein-coding genes: p=2.91*10^-4^, neurodegeneration screen hits: p=1.14*10^-5^). There is a significant difference in the slopes of the trends between age and gene expression for neurodegeneration screen hits and all protein-coding genes (all protein-coding genes: R=0.12, neurodegeneration screen hits: R=0.15, p=7.38*10^-6^). Regression lines indicate the relationship between age and TPM with a 95% confidence interval (standard error of the mean). The mixed effects regression analysis controlled for post-mortem interval, sex, ethnicity, and tissue of origin. b) Gene set enrichment plot showing that the set of age-associated neurodegeneration genes has reduced expression with respect to age. Vertical lines indicate rank of neurodegeneration screen hits by their association between gene expression and age determined by mixed-effects regression analysis coefficients. c) Proportion of genes that have significant associations between gene expression and age relative to the set of all protein-coding genes (blue) or the set of age-associated neurodegeneration genes (orange). Error bars indicate 95% binomial confidence intervals of the estimated proportion of genes with a significant association with age. Asterisk indicates tissues with an FDR-adjusted one-tailed hypergeometric test p-value less than 0.01. d) Proportion of protein-coding genes (blue) and age-associated neurodegeneration genes (orange) that are differentially expressed between Alzheimer’s disease (AD) and control in excitatory neurons in single-nucleus RNA-seq. Error bars indicate 95% binomial confidence intervals.

Next, we examined expression of the human orthologs of the screen hits with respect to age across regions of the human brain (Fig. 2c). Tissues enriched in age-associated changes of the screen hits include Alzheimer’s disease-vulnerable regions such as the hippocampus and the frontal cortex (Fig. 2c, hypergeometric test Benjamini-Hochberg FDR-adjusted p-value<0.1). In many cases, the same genes showed significant age-associated changes in expression in several tissues (Extended Data Fig. 1, mixed effect model Benjamini-Hochberg FDR-adjusted p-value<0.1, absolute value of regression coefficient>0.1). We observed that the Alzheimer’s disease-vulnerable tissues clustered together and with the Parkinson’s disease-vulnerable substantia nigra by hierarchical clustering (Extended Data Fig. 1). These human results suggest that the hits from our screen are associated with human aging in multiple regions of the brain, some of which are affected by common neurodegenerative diseases.

We analyzed the single nuclear RNA-seq data of excitatory neurons from a previously published single-nucleus RNA-seq study to examine cellular specificity of the neurodegeneration screen hits^66^. We observed that the average expression of screen hits was lower in Alzheimer’s disease-associated excitatory neurons than in excitatory neurons from healthy controls (Extended Data Fig. 2). We also found that the genes differentially expressed in Alzheimer’s disease-associated excitatory neurons in this dataset were enriched for neurodegeneration screen hits (Fig. 2d, Benjamini-Hochberg FDR-adjusted p-value<0.1). These results show that gene expression of the neurodegenerative screen hits declines with respect to age in human brain tissues and human Alzheimer’s disease excitatory neurons, suggesting their importance in human disease and aging.

### Human genetic risk factors enriched in disease-associated neurons complement results from the neurodegeneration screen

We collected human gene expression, human genetics, and proteomics, phosphoproteomics and metabolomics from *Drosophila* models of Alzheimer’s disease to systematically characterize omic perturbations in Alzheimer’s disease and further explore the role of neurodegeneration screen hits to human neurodegeneration (Fig. 1). For the human data, we used laser-capture microdissection to obtain pyramidal neurons from the human temporal cortex of 75 individuals, including 42 Alzheimer’s disease and 33 control individuals (Fig. 3a). We examined temporal pyramidal neurons because they are preferentially vulnerable to the formation of neurofibrillary tangles and neurodegeneration^35^. We performed RNA-seq and eQTL analysis on laser-captured material to examine how the hits from our transgenic *Drosophila* RNAi screen relate to genetic causes of Alzheimer’s disease in human neurons (Fig. 3a, TCPY in Supplementary Table 2). The RNA-seq from these pyramidal neurons demonstrated high expression of the excitatory neuron marker *SLC17A7*, as expected (Extended Data Fig. 3a)^67^.

**Fig. 3:**
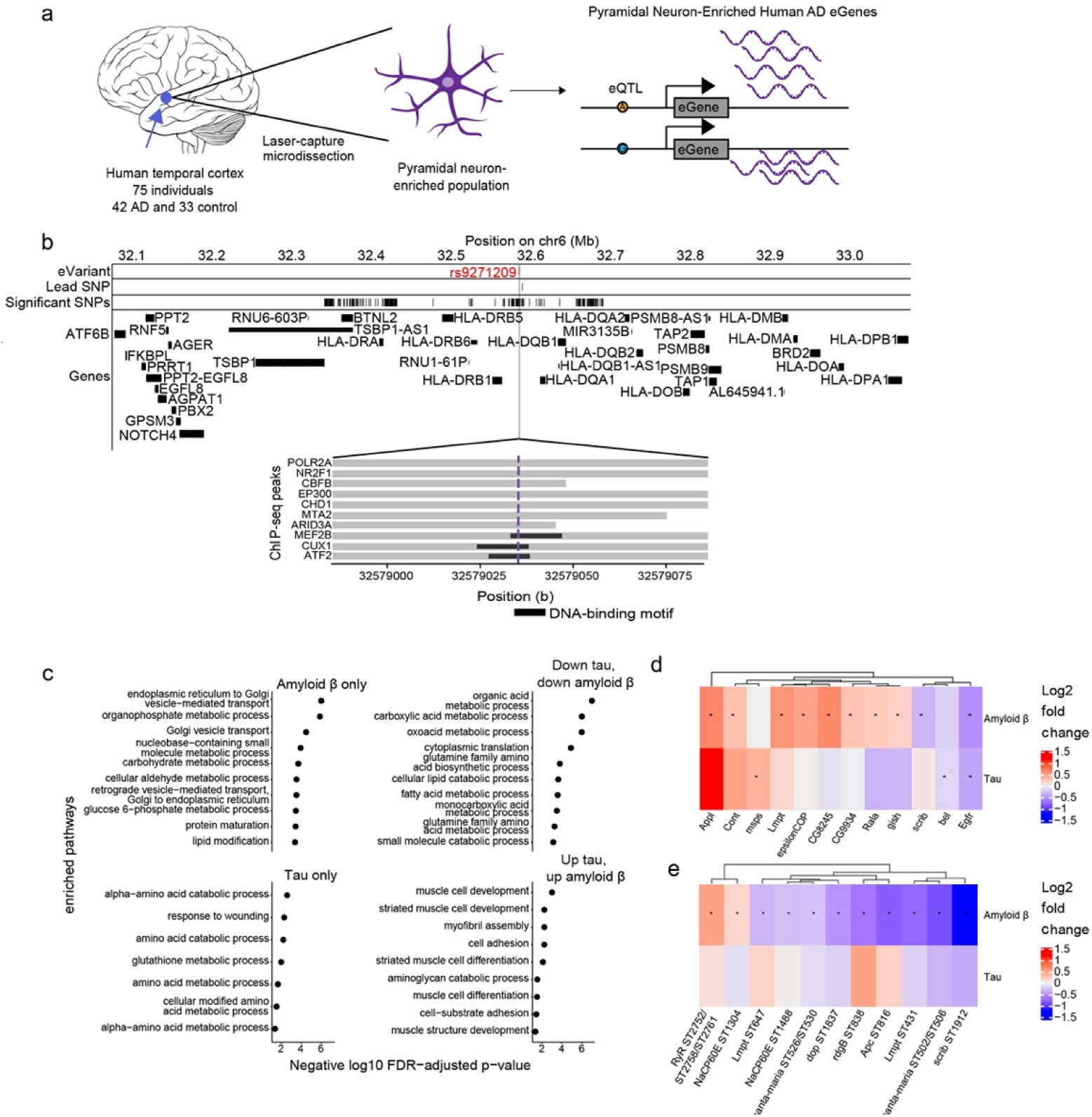
Multi-omic changes in human Alzheimer’s disease patients and model systems. a) Schematic depicting the identification of eGenes from laser-capture microdissection of temporal cortex pyramidal neuron-enriched populations from 75 individuals including 42 human Alzheimer’s disease (AD) and 33 healthy control patients and identification of eGenes. Brain cartoon created with Biorender.com. b) The eQTL associated with the eGene HLA-DRB1 is highlighted in red and overlaps with DNA binding motifs of MEF2B, CUX1 and ATF2 derived from ENCODE ChIP-seq and FIMO-detected motifs. Grey horizontal bars indicate ChIP-seq binding regions and the black horizontal bars indicate where the DNA-binding motif is located. c) Dot plots showing the negative log_10_ FDR-adjusted p-values for enriched GO terms in proteins that are significantly upregulated or significantly downregulated in both *Drosophila* models of tau and amyloid β, only differentially abundant in *Drosophila* models of amyloid β (Amyloid β only), or only differentially abundant in *Drosophila* models of tau (Tau only). d) Heat maps depict the log2 fold changes between Aβ_1-42_ transgenic flies (Amyloid β) or tau^R406W^ (Tau) transgenic flies with controls for d) proteins or e) phosphoproteins that were hits in the age-associated neurodegeneration screen. An asterisk indicates whether the comparison was significant at an FDR threshold of 0.1. The columns of all heatmaps were clustered by hierarchical clustering.

We first performed an eQTL meta-analysis across 7 different bulk RNA-seq and genomics studies in post-mortem brains (Supplementary Tables 2 and 3, Methods). The results from this meta-analysis were then forwarded to the eQTL analysis in the newly collected temporal cortex pyramidal neuron RNA-seq data to see which brain eQTLs were enriched in Alzheimer’s disease-vulnerable neurons. We found *cis-*regulatory effects in the pyramidal neuron-enriched transcriptomes for 12 eGenes (Table 2). The enriched genes included *C4A*, *EPHX2*, *PRSS36*, and multiple MHC class II genes (Table 2). Expression of the eGenes was correlated with several known biological processes previously associated with Alzheimer’s disease such as insulin signaling, protein folding and lipid metabolism^66,68–79^ (Extended Data Fig. 3b). We incorporated the eGenes from the temporal cortex pyramidal neurons and the meta-analysis in our analysis of the fly screen hits.

**Table 2.**
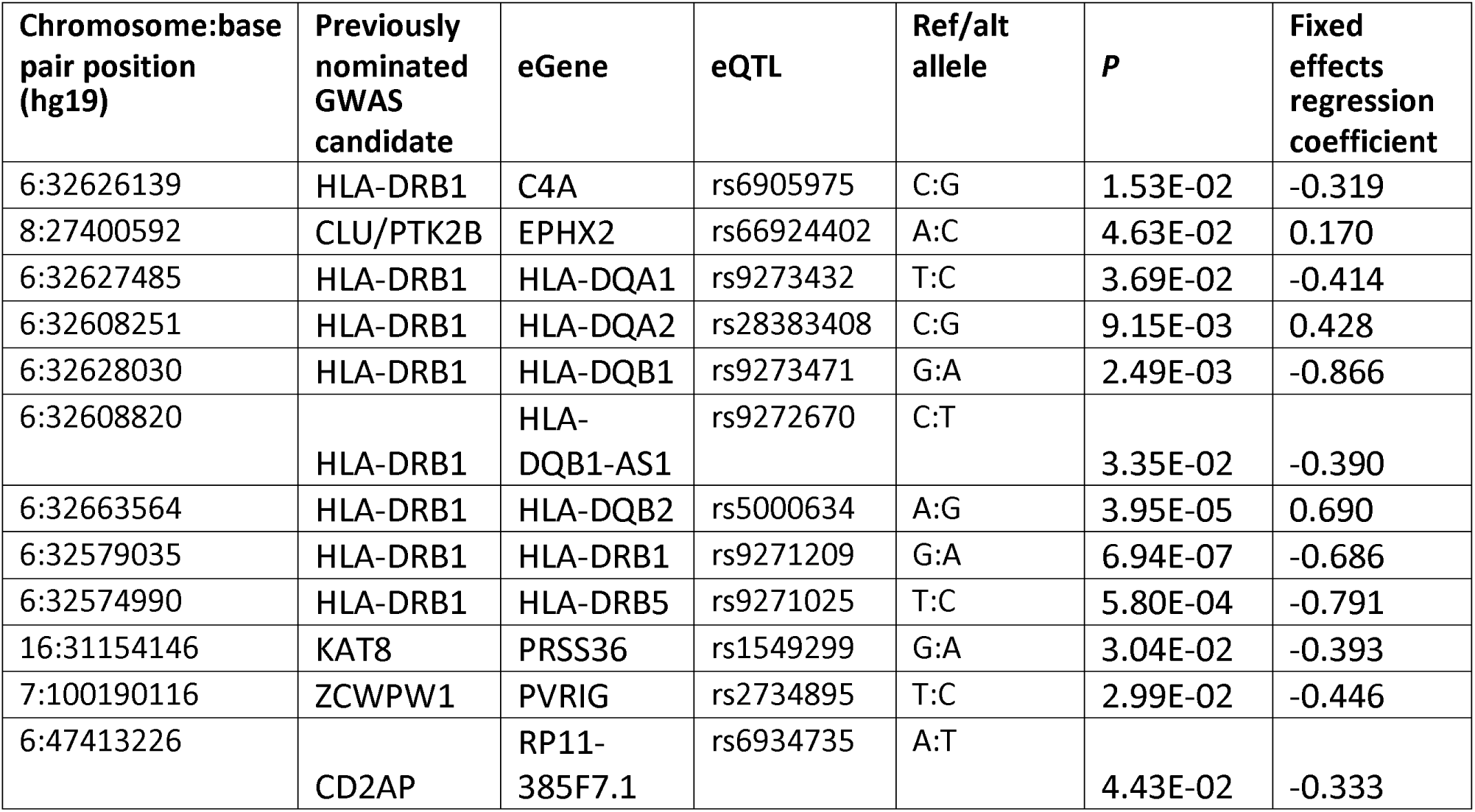
eQTLs linked to Alzheimer’s disease GWAS loci: eGenes and variants from an eQTL analysis of temporal cortex pyramidal neuron-enriched population from human Alzheimer’s disease and control patients. P-value from meta-analysis across 1087 human Alzheimer’s disease and control patients across 7 previously published studies is also reported. Beta coefficient indicates the association between gene expression of the eGene and presence of Alzheimer’s disease. Chromosomal coordinates are reported according to the human genome reference hg19 and the hypothetical gene is the variant reported in Jansen et al. 2019 for that particular locus^11^.

We hypothesized that some temporal cortex pyramidal neuron eQTLs influence eGene expression by disrupting transcription factor binding. We used the ENCODE 3 transcription factor ChIP-seq data to see which eQTLs overlapped transcription factor peaks and DNA-binding motifs (Fig. 3b). We found that the eQTL (*rs9271209*) for *HLA-DRB1* overlapped with ChIP-seq peaks and DNA-binding motifs for the transcription factors MEF2B, CUX1 and ATF2 (Fig. 3b). Patients with the *rs9271209* eQTL have reduced expression of *HLA-DRB1*, suggesting that this Alzheimer’s disease-associated effect on gene expression could be mediated through inhibition of transcription factor binding (Fig. 3b and Table 2).

### Proteomics, phosphoproteomics and metabolomics from Drosophila models of tauopathy or amyloid β neurotoxicity show significant changes in neurodegeneration screen hits in disease contexts

We generated proteomic, phosphoproteomic and metabolomic data from the heads of established *Drosophila* models of amyloid β and tau toxicity to broadly characterize molecular changes in models of Alzheimer’s disease pathology^80,81^ (Fig. 1 and Supplementary Tables 4-6). Specifically, we modeled amyloid β pathology using a transgenic fly line expressing the human amyloid β_1-42_ isoform (Aβ_1-42_ transgenic flies)^81^. We modeled tau pathology using a well-characterized transgenic fly line expressing human *MAPT* with the neurodegenerative disease-associated R406W point mutation (tau^R406W^ transgenic flies)^80^. We used tau^R406W^ transgenic flies because these flies display a modest, but detectable degree of neurodegeneration at 10 days of age^80^. We aged control and experimental flies for 10 days and measured proteomics, phosphoproteomics and metabolomics. Proteins downregulated in both Aβ_1-42_ transgenic flies and tau^R406W^ transgenic flies were enriched for enzymes that metabolize carboxylic acids, amino acids, and lipids, suggesting shared manifestations of molecular pathology (Fig. 3c, Benjamini-Hochberg FDR-adjusted p-value<0.1). Additionally, we found that proteins that were upregulated in Aβ_1-42_ transgenic flies and tau^R406W^ transgenic flies were enriched for muscle development and cell adhesion (Fig. 3c, g:SCS FDR-adjusted p-value<0.1). Proteins that were only differentially abundant in Aβ_1-42_ transgenic flies were enriched for Golgi-associated processes, while proteins that were only differentially abundant in tau^R406W^ transgenic flies were enriched for wound healing responses and amino acid synthesis pathways (Fig. 3c, Benjamini-Hochberg FDR-adjusted p-value<0.1). These results suggest biological processes that are common among models of Alzheimer’s disease, which are specific to amyloid β pathology, and which are specific to tau-mediated pathology.

Few of the neurodegeneration screen hits were differentially abundant in the proteomic and phosphoproteomic data, and the lack of overlap is a phenomenon that has been noted previously in other comparisons of genetic and expression data^82^. The screen hits that were differentially abundant in the Aβ_1-42_ transgenic fly proteomics were enriched for biological processes pertaining to development and cognition (Fig. 3d, Extended Data Fig. 4g, Benjamini-Hochberg FDR-adjusted p-value<0.1). None of the screen hits were differentially phosphorylated in the tau^R406W^ transgenic flies, while there were 11 phosphopeptides found in neurodegeneration screen hits that were differentially phosphorylated in Aβ_1-42_ transgenic flies (Fig. 3e). Despite the small overlap, our results highlight neurodegeneration screen hits that are differentially altered in models of Alzheimer’s disease pathology. Furthermore, we observe that more neurodegeneration screen hits show proteomic and phosphoproteomic changes in Aβ_1-42_ transgenic flies than in tau^R406W^ transgenic flies.

### Network integration of Alzheimer’s disease omics and novel genetic screening data identifies subnetworks representing biological processes underlying neurodegeneration in Alzheimer’s disease

We performed network integration of our *Drosophila* neurodegeneration screen hits with Alzheimer’s disease multi-omics to determine how the neurodegeneration screen hits contribute to human Alzheimer’s disease (Fig. 1). We integrated the hits from the neurodegeneration screen with our human eGenes and *Drosophila* proteomics, phosphoproteomics and metabolomics, a previously published genome-scale screen for tau mediated neurotoxicity tau^R406W^ flies, previously published human Alzheimer’s disease proteomics, and previously published human lipidomics using the Prize-collecting Steiner Forest algorithm (PCSF) to build a protein-protein/protein-metabolite interaction network model of Alzheimer’s disease^15,40,39^ (Figs. 1 and 4a; Supplementary Table 7). Briefly, the method links as many input nodes as possible while minimizing the number of low-confidence edges in the final network. The result includes a mix of the input nodes and new interactors of interest that were not part of the input set, which we call “predicted nodes” (Fig. 4a, Methods). We filtered out predicted nodes that did not show up in a sufficient number of networks after 100 edge permutations and were found in too many networks with randomized node weights (Supplementary Fig. 1, Methods). We found that the majority of the nodes in our network were observed in multiple hyperparameter selections, showing that these results were robust to parameter choices (Supplementary Table 7). This procedure filtered out 19102 nodes from the initial solution. We compared our results to a previously published study in a different disease indication, medulloblastoma, to emphasize the specificity of our results to Alzheimer’s disease. We found that while both analyses had subnetworks enriched for cell cycle regulation, none of the nodes in these overlapping enrichments were shared. This result supported our claim that our Alzheimer’s disease-associated network was distinct from networks derived from the same method in different disease indications^83^. The detailed results of this network are visualized in an interactive website (Methods, Data availability).

**Fig. 4:**
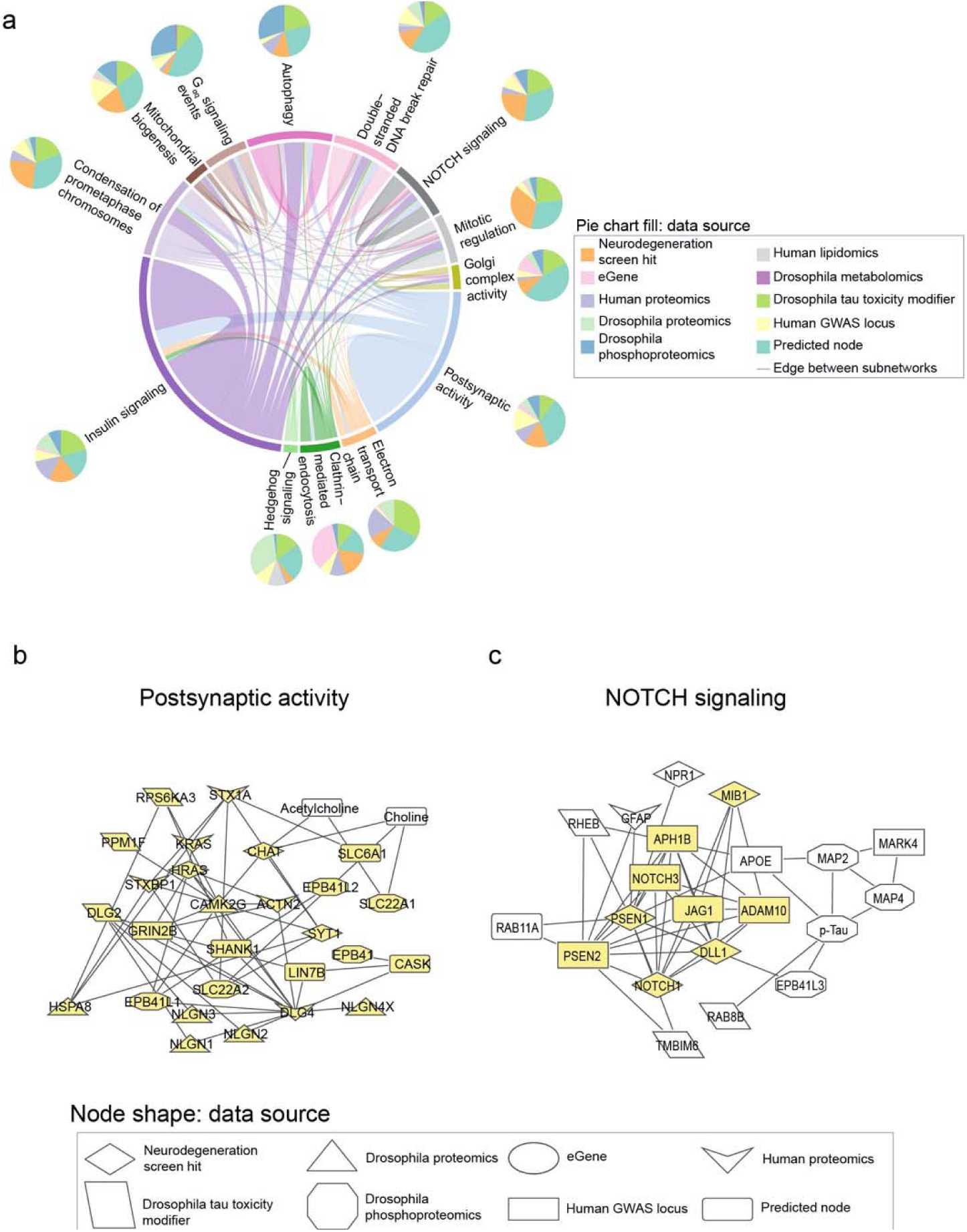
Network integration of Alzheimer’s disease multi-omics and novel genetic screening data identifies subnetworks characterized by hallmarks of neurodegeneration and processes previously not implicated in Alzheimer’s disease. a) Network integration of human and *Drosophila* multi-omics for Alzheimer’s Disease highlights subnetworks enriched for proteins belonging to known gene ontologies. Each subnetwork is represented by a pie chart, which indicates the proportion of nodes represented by a given data type. Edge width is determined by the number of interactions between nodes within or with another subnetwork and colored by one of the involved subnetworks. Each pie chart is labeled by the enriched biological process by hypergeometric test (FDR-adjusted p-value less than 0.1). b) A subnetwork enriched for postsynaptic activity. Nodes belonging to the annotated process are highlighted in yellow. Also in this subnetwork are metabolites associated with postsynaptic activity such as acetylcholine. c) Phosphorylated tau, APOE, and APP-processing proteins interact with each other and are in a subnetwork enriched for NOTCH signaling-associated genes. Members of the NOTCH signaling pathway are highlighted in yellow.

Louvain clustering of the network revealed subnetworks enriched for biological processes associated with Alzheimer’s disease in previous studies, such as insulin signaling, postsynaptic activity, and double-strand break repair^71–73,78,79,84–86^ (Fig. 4a). Subnetworks were also enriched for cell signaling pathways such as NOTCH signaling and hedgehog signaling that have not been previously characterized as hallmarks of neurodegeneration^79^ (Fig. 4a). We chose Louvain clustering instead of Leiden clustering because the Louvain clusters had a higher modularity score than the Leiden clusters (Louvain Q=0.415, Leiden Q=0.412). We note that there is a significant overlap in the number of biological processes enriched in the subnetworks derived from Louvain clustering or Leiden clustering, suggesting that our findings are robust to the clustering method of choice (hypergeometric test p-value<1*10^-16^).

506 of the 1008 nodes in the network had no previously known association with Alzheimer’s disease according to OpenTargets, which supports the ability of the network-based approach to identify new regulators of biological processes in Alzheimer’s disease. Out of the 690 input nodes in the network, only 60 were enriched in more than one data type. The largest intersection was of 15 nodes shared between *Drosophila* amyloid β proteomics and phosphoproteomics. These overlapping nodes were enriched for ERBB4 signaling, NRIF-mediated cell death and ion transport (Supplementary Table 7). These observations highlight the contribution of network analysis in integrating findings from multiple data sources to discover disease-associated biological processes and to design experiments about specific nodes of interest.

We re-ran the Prize-Collecting Steiner Forest omitting one data type at a time to determine the relative contribution of each data type to the network solution. Most of the nodes that were missing from these new network solutions were those contributed by the omitted data type (Supplementary Table 7). Regardless of the data type removed, each PCSF result had a subnetwork enriched for mitochondrial biogenesis and at least one pathway involved in postsynaptic activity (Supplementary Table 7). We note that the networks that lacked neurodegeneration screen hits were not enriched for hedgehog signaling, insulin signaling, autophagy, Gɑq signaling events, and condensation of prometaphase chromosomes like our final network (Supplementary Table 7). Overall, removing the neurodegeneration screen hits removed the largest number of pathway enrichments compared to the final network solution (Supplementary Table 7). In contrast, removing Alzheimer’s disease GWAS hits removed pathway enrichments for Gαq signaling events and NOTCH signaling (Supplementary Table 7). These results underscore the importance of neurodegeneration screen hits in our network analysis, and how each integrated datatype influenced the enrichment of different biological processes in our final network solution.

We inspected the nodes of our network communities to determine whether the subnetworks represented expected or new relationships in the context of Alzheimer’s disease. The subnetwork enriched for postsynaptic activity showed expected protein-metabolite and protein-protein interactions in choline metabolism^87,88^ (Fig. 4b). We observed interactions involving the metabolite acetylcholine with choline O-acetyl transferase (CHAT) and choline transporter (SLC22A1) (Fig. 4b). Additionally, we saw interactions between choline, CHAT and choline transporters SLC22A1 and SLC22A2 (Fig. 4b). This subnetwork illustrates the ability of our network analysis to recover established biological processes in Alzheimer’s disease.

A novel role of NOTCH signaling emerged in one subnetwork that linking members of the pathway with phosphorylated tau, members of the gamma secretase complex, the APOE protein (Fig. 4c). Each of these proteins has been associated with hallmarks of Alzheimer’s disease^89–93^. This subnetwork suggests that Notch signaling could influence the previously characterized relationship between amyloid β and APOE^94^. These results suggest roles for NOTCH signaling proteins in Alzheimer’s disease-mediated pathology.

We found strong overlaps for pathway enrichments between our network and networks derived from comparable methods. We compared our result to those of ROBUST and GNNExplainer (Supplementary Table 7). We found a significant overlap in the number of enriched biological pathways in subnetworks derived from the Prize-Collecting Steiner Forest and ROBUST (hypergeometric p-value<1*10^-16^). We further found a significant overlap between nodes in the Prize-Collecting Steiner Forest-derived network and the ROBUST-derived network (hypergeometric p-value<1*10^-16^). Some pathways were only enriched in one method: viral response pathways were only enriched in the ROBUST-derived network, inositol phosphate metabolism pathways were only enriched in the GNNExplainer-derived network and respiratory electron transport pathways were only enriched in the Prize-Collecting Steiner Forest results. Overall, our network analysis findings were not highly dependent on our network integration method of choice.

### Network integration of Alzheimer’s disease Omics and genetic hits reveals HNRNPA2B1 and MEPCE as targets that regulate tau-mediated neurotoxicity

We then experimentally tested implications of a subnetwork linking a screen hit (*HNRNPA2B1)* and an eGene (*MEPCE*) with *Drosophila* modifiers of tau toxicity^15^ (Fig. 5a). *HNRNPA2B1* is known as an intronic splice suppressor, while *MEPCE* is known as a protein that provides a methyl phosphate cap on 7SK non-coding RNA. *HNRNPA2B1* was of particular interest because it was the only neurodegeneration screen hit that interacted with phosphorylated tau in our network (Extended Data Fig. 5). We selected *MEPCE* for follow-up experimentation because it interacted with *HNRNPA2B1,* it did not previously have an association with tau-mediated neurotoxicity, and had the highest-confidence *Drosophila* ortholog among the hits from our eGene analysis. We knocked down the fly orthologs of *HNRNPA2B1* or *MEPCE* in a *Drosophila* model of tauopathy with two independent RNAi lines per gene (Figs. 5b,c). To increase relevance to Alzheimer’s disease, in which wild type human tau is deposited into neurofibrillary tangles, we used transgenic flies expressing wild-type human tau (tau^WT^) in the fly retina^80^. We found that knockdown of fly orthologs of either *HNRNPA2B1* or *MEPCE* enhanced tau retinal toxicity, as quantified using a previously described semi-quantitative rating scale based on morphologic disruption and reduction in size of the adult fly eye following expression of human tau^95^ (Figs. 5b,c, one-way ANOVA with Tukey’s post-hoc correction p<0.05). Enhancement of tau toxicity in the fly retina is consistent with the human eQTL results, which show that *MEPCE* expression is reduced in Alzheimer’s disease patients with the eQTL *rs7798226* (Supplementary Table 3) and suggest a mechanism for effects of the GWAS variant in Alzheimer’s disease.

**Fig. 5:**
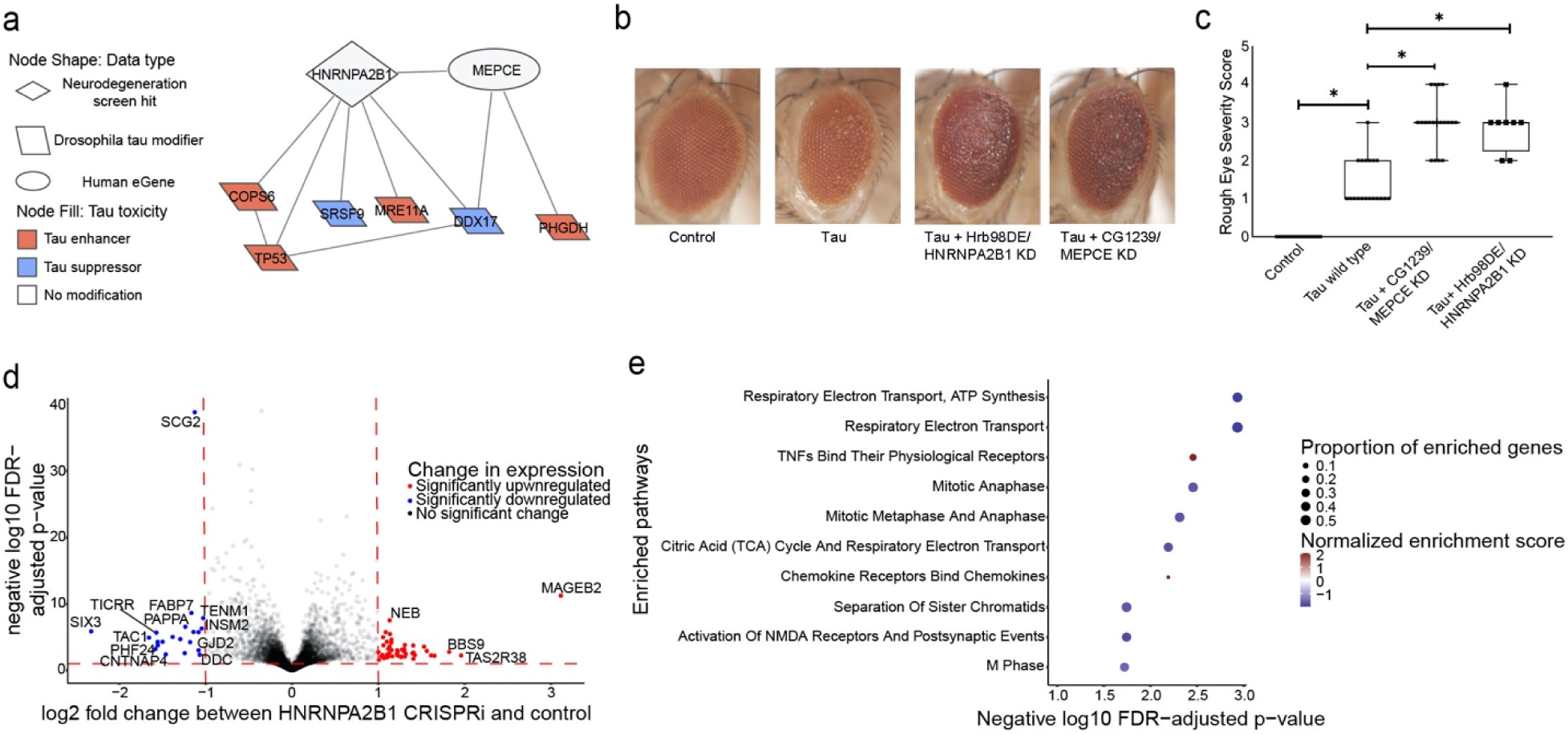
Network integration of Alzheimer’s disease multi-omics and novel genetic screening data reveals biological processes associated with tau-mediated neurotoxicity. a) The neurodegeneration modifier *HNRNPA2B1* and the eGene *MEPCE* interact with each other and have protein-protein interactions with modifiers of tau neurotoxicity. The interaction between HNRNPA2B1 and MEPCE is found in the subnetwork in Figure 4 that is enriched for insulin signaling. b) Knockdown of the *Drosophila* orthologs of *HNRNPA2B1* (*Hrb98DE*) and *MEPCE* (*CG1293*) shows enhancement of the rough eye phenotype in flies expressing wild type human tau. c) Quantification of rough eye severity. The scale reflects the extent of morphological disruption after human tau retinal expression (Methods). Statistical significance was measured using a one-way ANOVA with Tukey’s post-hoc correction and is indicated with an asterisk. Error bars are the standard error of the mean. Two independent RNAi constructs were used to knock down each gene. n=8. Control is *GMR-GAL4/+*. Flies are one day old. d) Volcano plot depicting differential expression analysis by DeSeq2 of bulk RNA-seq after *HNRNPA2B1* CRISPRi knockdown in NGN2 neural progenitor cells (Benjamini-Hochberg FDR<0.1, absolute log_2_ fold change > 1). Each dot represents a single gene. The horizontal dashed line indicates the negative log_10_ FDR-adjusted p-value significance cut-off of 0.1 and the vertical dashed lines indicate the log_2_ fold change cut-offs of 1 and -1. Red dots indicate genes that are significantly upregulated and blue dots indicate genes that are significantly downregulated. e) Dot plot of the enriched pathways identified by gene set enrichment analysis of the RNA-seq data. The 10 pathways with the highest negative log_10_ FDR-adjusted p-value are plotted. The size of the dot indicates the proportion of genes that are part of the enriched pathway. The color of the dot represents the normalized enrichment score (NES), where blue indicates downregulation and red indicates upregulation. The x-position of the dot indicates the negative log_10_ FDR-adjusted p-value and the y-position is the corresponding, enriched pathway.

To understand how *HNRNPA2B1* contributes to age-associated neurodegeneration in human systems, we performed RNA-seq after CRISPRi knockdown of *HNRNPA2B1* in human iPSC-derived, NGN2 neural progenitor cells. Our knockdown achieved a partial reduction of *HNRNPA2B1* gene relative to control (Extended Data Fig. 6a, log_2_(Fold Change)=-0.60, Benjamini-Hochberg FDR-adjusted p-value<0.1). Differential expression after *HNRNPA2B1* knockdown showed that the most significantly downregulated genes involved those involved in neuronal development or synaptic activity such as *SCG2*, *FABP7*, *TENM1*, and *SIX3* (Fig. 5d and Supplementary Table 8, log_2_(Fold Change)<-1, Benjamini-Hochberg FDR-adjusted p-value<0.1). No individual subnetwork was enriched for differentially expressed genes after *HNRNPA2B1* knockdown. Gene Set Enrichment Analysis showed that the top enriched pathways include downregulation of the electron transport chain and of genes involved in postsynaptic events (Fig. 5e and Supplementary Table 9, Benjamini-Hochberg FDR-adjusted p-value<0.1). Reduced postsynaptic activity and electron transport chain activity have been previously associated with Alzheimer’s disease and tau-mediated neurotoxicity^78,96–101^. These changes suggest potential mechanisms by which *HNRNPA2B1* contributes to tau-mediated neurotoxicity and neurodegeneration in human aging.

### Network analysis implicates CSNK2A1 and NOTCH1 as regulators of the Alzheimer’s disease-associated biological process of DNA damage repair in neurons

In addition to the network connections between NOTCH signaling proteins and hallmark proteins of Alzheimer’s disease (Fig. 4c), we also noted that NOTCH signaling proteins were linked with the Alzheimer’s disease-associated process of DNA damage, a process also associated with Alzheimer’s disease^67,84–86,102,103^(Fig. 6a). We calculated the network betweenness of all neurodegeneration screen hits with regulators of DNA damage repair (Supplementary Table 10). We found that *NOTCH1* and the neurodegeneration screen hit *CSNK2A1* were among the top 5 neurodegeneration genes in terms of network betweenness among nodes associated with DNA damage repair (Supplementary Table 10). CSNK2A1 and NOTCH1 shared some interactors that regulated DNA damage repair (Fig. 6a). NOTCH1 is a key receptor in NOTCH signaling that regulates cell-fate decisions. *CSNK2A1* is a kinase for many proteins including casein; the activity of *CSNK2A1* regulates processes such as apoptosis and cellular proliferation. All the interacting DNA damage repair-associated nodes that interact with *CSNK2A1* and *NOTCH1* except for *H2AFX* and *COPS2* regulate double-strand break repair, suggesting that *CSNK2A1* and *NOTCH1* knockdown may disrupt the DNA damage response.

**Fig. 6:**
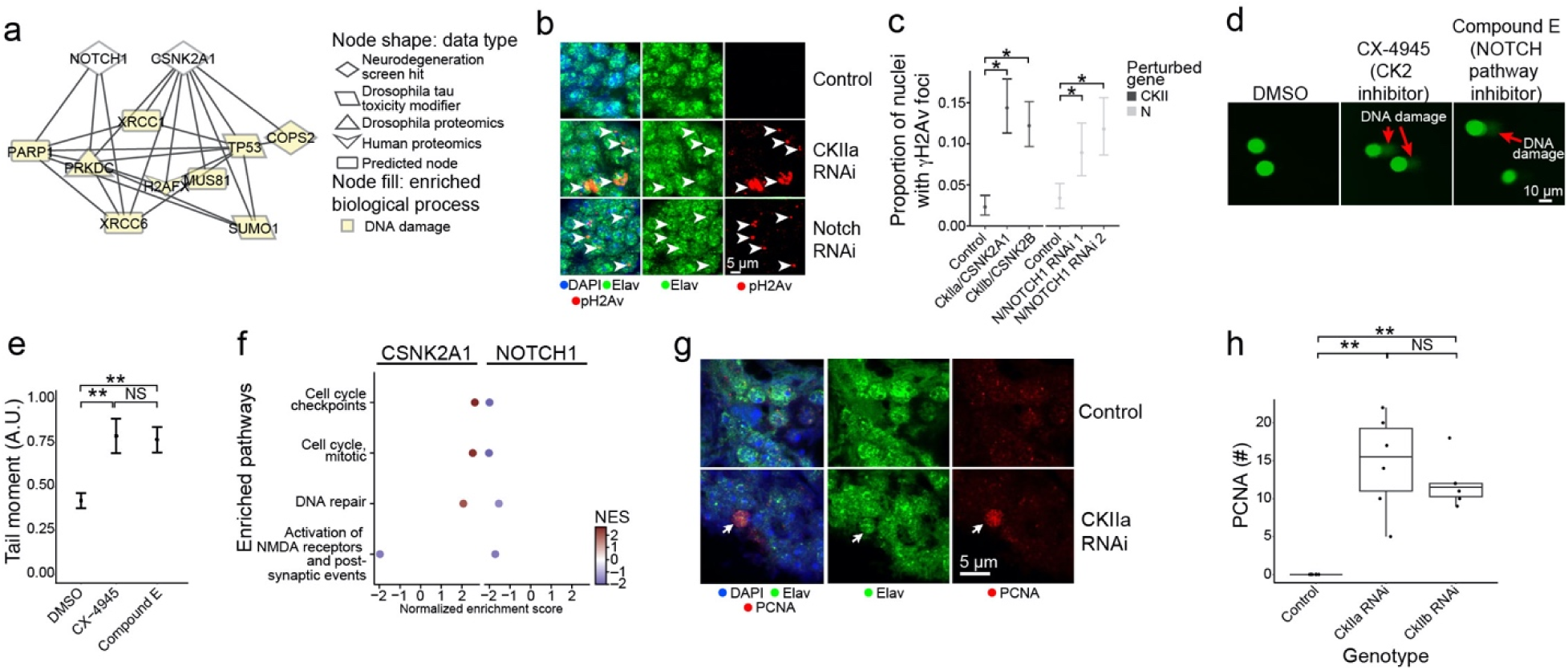
Network analysis implicates neurodegeneration genes as regulators of the AD-associated biological process of DNA damage repair. a) NOTCH1 and CSNK2A1 interact with a diversity of AD-specific omics that are involved in DNA damage repair processes. Nodes involved in DNA damage are highlighted in yellow. b) Knockdown of *Drosophila* orthologs for *NOTCH1* and *CSNK2A1* lead to increased DNA damage in neurons of the adult fly brain as measured by increased numbers of foci positive for the DNA double-strand break marker pH2Av (red, arrowheads). pH2Av is the *Drosophila* ortholog of mammalian pH2AX. Neurons are identified by elav immunostaining (green). Nuclei are identified with DAPI immunostaining (blue). The scale bar represents 5 µm. c) Percent of nuclei containing pH2Av foci in control flies, *Drosophila* knockdowns of orthologs of *CSNK2A1* (*CKIIa* and *CKIIb*) and *NOTCH1* (N). Asterisks indicate significance of a one-way binomial test after Benjamini-Hochberg FDR correction p<0.01. Error bars are 95% binomial confidence intervals. n=6. Controls are *elav-GAL4/+; UAS-Dcr-2/+* (CKII knockdown) or *elav-GAL4/+* (N knockdown). Flies are 10 days old. d) Inhibition of Casein Kinase 2 (CK2) by CX-4945, and the inhibition of NOTCH cleavage by Compound E enhances DNA damage in human iPSC-derived neural progenitor cells measured by the COMET assay. e) Quantification of the tail moments from panel A in arbitrary units. Asterisks indicate p<0.01 by ANOVA with Tukey’s Post-Hoc correction. Error bars indicate standard error of the mean. f) Dot plots showing the normalized enrichment scores (NES) of selected, significantly enriched REACTOME pathways after *CSNK2A1* and *NOTCH1* knockdown in NGN2 neural progenitor cells. Red and blue dots indicate positive and negative NES, respectively, reflecting upregulation or downregulation of pathways. Pathways were selected to show shared changes in pathways related to cell cycle, DNA repair and postsynaptic activity. g) Representative immunofluorescence images of mature neurons in *Drosophila* brains show inappropriate cell cycle re-entry in postmitotic neurons as indicated by PCNA expression (red, arrow) following *CKIIa* knockdown. The neuronal marker elav identifies neurons (shown in green). PCNA, elav (neurons) and DAPI are represented in red, green, and blue respectively. h) Quantification of PCNA expression in control brains and brains of *Drosophila* with knockdown of orthologs of *CSNK2A1* (*CKIIa* and *CKIIb*). Asterisks indicate p<0.01 by ANOVA with Tukey’s Post-Hoc correction. n=6. Control is *elav-GAL4/+; UAS-Dcr-2/+.* Flies are 10 days old.

Next, we used RNAi to knock down *Drosophila* orthologs of *NOTCH1* and *CSNK2A1* in a pan-neuronal pattern in the aging adult fly brain to assess the relationship between these neurodegeneration screen hits and DNA damage (Figs. 6b,c). To exclude off-target effects we used two independent RNAi transgenes targeting the *NOTCH1* ortholog N (Figs. 6b,c). For *CSNK2A1* we used one RNAi to target the *CkIIa* subunit of the casein kinase holoenzyme and another RNAi to target the *CkIIb* subunit of the casein kinase holoenzyme (Figs. 6b,c) because other available transgenic RNAi lines were lethal with pan-neuronal expression. We observed that knockdown of the *Drosophila* orthologs for *NOTCH1* and *CSNK2A1* led to an increase in DNA damage, as measured by the number of phospho-H2Av (*Drosophila* ortholog of mammalian phosphor-H2AX) foci (Figs. 6b, arrowheads, c, One-Way Binomial Test p<0.01).

We performed a COMET assay in wild-type human neuronal progenitor cells treated with inhibitors for the Notch signaling pathway or the casein kinase holoenzyme (CK2) to test if reduced *CSNK2A1* or *NOTCH1* function leads to increased DNA damage in human cells (Figs. 6d, arrows, e). We observed that treatment with the Notch inhibitor Compound E and the CK2 inhibitor CX-4945 led to an increase in the tail moment of the neural progenitor cells compared to DMSO treatment, showing an increase in DNA damage after inhibitor treatment (Fig. 6d, arrows, e, ANOVA with Tukey’s post-hoc correction p-value<0.01, Methods). These results show how the screen hits *NOTCH1* and *CSNK2A1* regulate DNA damage in human and *Drosophila* neurons, as inferred by our computational network analysis.

### Transcriptomic analysis suggests how CSNK2A1 and NOTCH1 knockdown could lead to age-associated neurodegeneration through distinct DNA-damaging pathways

We performed RNA-seq after CRISPRi knockdown of *CSNK2A1* or *NOTCH1* in NGN2-expressing neural progenitor cells to broadly understand how human cells respond to reduced *CSNK2A1* and *NOTCH1* expression (Fig. 6). Expression of both target genes dropped significantly in the respective knockdowns (*CSNK2A1*: log_2_(fold change)<-1, FDR-adjusted p-value<0.1, Extended Data 6b; *NOTCH1:* log_2_(fold change)=-0.92, FDR-adjusted p-value<0.1, Extended Data Fig. 6c), with good clustering of replicates in PCA analysis (Extended Data Fig. 6d). We found 145 significantly upregulated and 282 significantly downregulated genes upon knocking down *CSNK2A1*, while we found 15 significantly upregulated and 5 significantly downregulated genes after knocking down *NOTCH1* (Extended Data Fig. 7a,b and Supplementary Table 8, absolute value of log_2_(fold change)>1, FDR-adjusted p-value<0.1). The disparity in the number of differentially expressed genes could be explained by how the knockdown efficiency of *NOTCH1* was less than that of *CSNK2A1* (Extended Data Fig. 6a,b). We then used Gene Set Enrichment Analysis (GSEA), which can identify coordinated expression changes, even when some fall below univariate statistical thresholds. GSEA found significant enrichment for DNA damage repair pathways. However, we were surprised to find that these pathways were upregulated after *CSNK2A1* knockdown but downregulated after *NOTCH1* knockdown (Fig. 6f and Extended Data Table 7). Interestingly, we also found an upregulation for cell cycle regulating-genes after *CSNK2A1* knockdown (Fig. 6f).

Examining the expression changes in the context of the network, we found that that the effects of *CSNK2A1* knockdown decreased significantly the greater the network distance from *CSNK2A1* (Kruskal-Wallis test p=4.77*10^-4^, Extended Data Fig. 7c). There was no significant variation in log_2_ change between *NOTCH1* knockdown and control with respect to the degree of separation from *NOTCH1*, which could also be a consequence of lower *NOTCH1* knockdown efficiency than *CSNK2A1* knockdown efficiency (Kruskal-Wallis test p=0.146, Extended Data Fig. 7d).

To determine if *CSNK2A1* knockdown could alter cell cycle in aging postmitotic neurons in vivo, we knocked down the *Drosophila* the *CkIIa* and *CKIIb* subunits of the casein kinase holoenzyme in adult *Drosophila* brain neurons and assessed changes in proliferating cell nuclear antigen (PCNA), a robust marker of cell cycle activation in *Drosophila* and mammalian systems^20,104,105^ (Fig. 6g, arrow, h). We found a significant increase in PCNA following *CKIIa* knockdown of either *CKIIa* or *CKIIb* (Fig. 6h, ANOVA with Tukey’s post-hoc correction p=4.94*10^-5^), supporting our hypothesis that knockdown of *CSNK2A1* promotes neuronal activation of cell cycle regulators. As expected, there was no PCNA activation in control postmitotic neurons (Fig. 6g,h). Activation of cell cycle proteins in mature neurons is associated with Alzheimer’s disease, cell death, and double-strand break-bearing neurons^20,106–109^. Our results collectively suggest that *CSNK2A1* knockdown may lead to neurodegeneration through accumulation of DNA damage and subsequent inappropriate activation of the cell cycle in postmitotic neurons leading to cell death.

## Discussion

We have integrated results from the largest reported forward genetic screen for age-related neurodegeneration mutants with multi-omic data from Alzheimer’s disease patients and Alzheimer’s disease relevant models to computationally and experimentally define pathways controlling neurodegeneration. One highlight of our work is the demonstration that *CSNK2A1* and *NOTCH1* regulate age-associated neurodegeneration through DNA damage response pathways (Figs. 5 and 6). We provide evidence that *CSNK2A1* and *NOTCH1* regulation of the Alzheimer’s disease-associated process of DNA damage is a key process for neurodegeneration among the many processes these genes regulate. Previous studies showed that that HDAC inhibitors reduced DNA damage burden and neuronal cell death^67,84,102,103,110–112^. Other studies have proposed neuroprotective compounds that inhibit cell cycle re-entry in postmitotic neurons like we observed upon *CSNK2A1* knockdown^113^. Our study suggests that the *CSNK2A1* and *NOTCH1* pathways should also be explored for potential approaches to prevent DNA damage-associated neurodegeneration.

Future work could explore cause-and-effect relationships between DNA damage and activation of cell cycle genes in the context of *CSNK2A1* knockdown. Currently, the relationship between cell cycle regulators and DNA damage in neurodegeneration is unclear^114^. One hypothesis supported by our results is that *CSNK2A1* knockdown leads to neurodegeneration by activating genes that promote DNA replication and entry into the G_1_ phase of the cell cycle, amplifying existing DNA damage in the neuron (Figs. 6d,e). Alternatively, excess accumulation of DNA damage upon *CSNK2A1* knockdown could lead to inappropriate activation of cell cycle regulators and DNA repair proteins to fix DNA damage (Figs 6d,e). Follow-up work can determine the causes or consequences of DNA damage as regulated by *CSNK2A1*. Such studies can help inform neuroprotective approaches for limiting age-associated DNA damage.

In another advance from our study, we suggest how changes in *MEPCE* expression contribute to neuronal death in Alzheimer’s disease (Fig. 5). Our eQTL analysis showed that patients that inherited the *rs7798226* eQTL had reduced *MEPCE* expression and our experimental data shows that reduced expression of *MEPCE* enhances tau toxicity in the *Drosophila* nervous system (Figs. 5b,c and Supplementary Table 2). Future studies could investigate whether the downregulation of *MEPCE* in patients with the *rs7798226* eQTL is strong enough to induce tau-mediated neurotoxicity in humans. This example illustrates how multi-omic network integration identified pathways potentially downstream of a disease-causing variant. Our network analysis work identified an eQTL that may play a role in Alzheimer’s disease-mediated neurodegeneration, which is an inference that could not be made from fine-mapping analysis alone.

Interestingly, some of our network findings differ from expectations in the literature. We found from our network analysis and subsequent experimentation in human tau transgenic flies that knockdown of *HNRNPA2B1* led to increased age-associated neurodegeneration and increased tau-mediated neurotoxicity (Fig. 5). However, *HNRNPA2B1* was upregulated in Alzheimer’s disease excitatory neurons in the largest published single nucleus RNA-seq study in human Alzheimer’s disease^67^. Another study showed that *HNRNPA2B1* knockdown in iPSC-derived neurons and mouse hippocampal neurons was protective against oligomeric tau-mediated neurotoxicity^115^. Given these prior observations, our results suggest that the *HNRNPA2B1* is under tight control; significant changes in *HNRNPA2B1* homeostasis may have consequences on tauopathy.

The network algorithms used in this study have previously been used to identify biological processes in various disease consequences, including Alexander disease, medulloblastoma, Parkinson’s disease in *Drosophila*, amyotrophic lateral sclerosis, and an *Appl* model of Alzheimer’s disease in *Drosophila*^28,83,116–118^. Related algorithms such as ROBUST, COSMOS, DOMINO and GNN-based tools have demonstrated the broad applicability of such approaches^119–122^. In some cases, the goal of network integration is to identify novel targets that were not found in the input data, but are implicated by the network. Indeed, in prior work, we knocked down a large number of nodes nominated by the network to test for neurotoxicity in a *Drosophila* model relevant to Amyotrophic Lateral Sclerosis and found that many of these targets had a significant effect on neurotoxicity^118^. The goal of the current study was different. Our genetic screen was intentionally broad, with genes knocked down in a pan-neuronal pattern to maximize recovery of neurodegeneration hits. The comprehensiveness of the screen reduced the need for computational methods to expand the number of genes of interest. Rather, the role of computational was to prioritize the genes and to generate hypotheses explaining the mechanisms by which these genes influenced neuronal health.

Many of the nodes in our network had only modest effect sizes in comparisons between disease in control. Additionally, the genes we studied in our follow-up experiments had only modest prior connections with Alzheimer’s disease; in the OpenTargets list of Alzheimer’s associated genes, the highest rank was 1256^th^ out of 6595. The network-based multi-omic integration allowed us to focus on disease-associated targets that would not be found by simpler approaches.

The framework presented in this paper could be used to combine the screen hits with appropriate disease-specific data to search for disease-universal or disease-specific regulators across neurodegenerative diseases. We observed that a significant proportion of age-associated genes in multiple human brain tissues are enriched for neurodegeneration screen hits. Given the diversity of brain regions affected in aging-related disorders, some of the screen hits are likely associated with diseases other than Alzheimer’s disease, and some may influence more than one disease. We also note that while our genetic screen data was neuron-specific, future work could use network analysis approaches presented in this or other studies to screens in other non-neuronal cell types^10,123^. Pathways that influence multiple diseases would be particularly important for therapeutic strategies to prevent aging of the brain.

## Methods

### Data availability

The full network is available and explorable at https://fraenkel-lab.github.io/neurodegeneration-network/

### Code availability

Code can be found at https://github.com/fraenkel-lab/neurodegeneration-network

#### Drosophila *stocks and Genetics*

All fly crosses and aging were performed at 25°C. Equal numbers of adult male and female flies were analyzed. For the genome-scale screen, gene knockdown was mediated by the *elav-GAL4* pan-neuronal driver and brain histology was examined at 30 days post-eclosion. Transgenic RNAi lines for genome-scale gene were obtained from the Bloomington *Drosophila* Stock Center and from the Vienna *Drosophila* Resource Center and are listed in Supplemental Table 1. We used all available transgenic RNAi lines from the Bloomington *Drosophila* stock center when the screen was performed. The *UAS-tau wild type, UAS-tau^R406W^ and UAS-Aβ^1-42^* transgenic flies been described previously^80,81^. Expression of human tau or amyloid β was directed to neurons using the pan-neuronal driver *elav-GAL4* or to the retina using the *GMR-GAL4* driver. Flies were aged to 10 days post-eclosion for brain proteomics, metabolomics, and histology. Dcr-2 was expressed in some animals to enhance RNAi-mediated gene knockdown. The following stocks were also obtained from the Bloomington *Drosophila* Stock Center: *elav-GAL4, GMR-GAL4, UAS-CG1239 (MEPCE) RNAi HMC02896, UAS-CG1239 (MEPCE) RNAi HMC04088, UAS-Hrb98DE (HNRNPA2B1) RNAi HMC00342, UAS-Hrb98DE (HNRNPA2B1) RNAi JF01249, UAS-CkIIɑ RNAi JF01436, UAS-CkIIβ RNAi JF01195, UAS-N RNAi 1 (GLV21004), UAS-N RNAi 2 (GL0092), UAS-Dcr-2*.

#### Histology, immunostaining, and imaging

Flies were fixed in formalin and embedded in paraffin. 4 µm serial frontal sections were prepared through the entire brain and placed on a single glass slide. Hematoxylin and eosin staining was performed on paraffin sections. For immunostaining of paraffin sections, slides were processed through xylene, ethanol, and into water. Antigen retrieval by boiling in sodium citrate, pH 6.0, was performed prior to blocking. Blocking was performed in PBS containing 0.3% Triton X-100 and 2% milk for 1 hour and followed by incubation with appropriate primary antibodies overnight. Primary antibodies to the following proteins were used at the indicated concentrations: pH2Av (Rockland, 600-401-914, 1:100), elav (Developmental Studies Hybridoma Bank, 9F8A9 at 1:20 and 7E8A10 at 1:5) and PCNA (DAKO, MO879, 1:500). For immunofluorescence studies, Alexa 555-and Alexa 488-conjugated secondary antibodies (Invitrogen) were used at 1:200. For quantification of pH2Av, a region of interest comprised of approximately 100 Kenyon neurons was identified in well-oriented sections of the mushroom body and the number of neurons containing one or more than one immuno-positive foci was determined. Images were taken on Zeiss LSM800 confocal microscope (Carl Zeiss, AG), and quantification was performed using Image-J software. All acquisition parameters were kept the same for all experimental groups. Quantification for PCNA was assessed by counting the number of sections containing PCNA immunopositivity in the entire brain. At least 6 brains were analyzed per genotype and time point for pH2Av and PCNA quantification. Histologic assessments were performed blinded to genotype.

#### Quantitative Mass Spectrometry sample preparation for proteomics

Three control (genotype: *elav-GAL4/+),* three tau (genotype: *elav-GAL4/+; UAS-tau^R406W^/+),* and three Aβ_1-42_ (genotype: *elav-GAL4/+; UAS-Aβ^1-42^)* samples of approximately 350 fly heads each were used for proteomic analysis. Samples were prepared as previously described (Paulo and Gygi 2018) with the following modifications. All solutions are reported as final concentrations. *Drosophila* heads were lysed by sonication and passaged through a 21-gauge needle in 8 M urea, 200 mM EPPS, pH 8.0, with protease and phosphatase inhibitors (Roche). Protein concentration was determined with a micro-BCA assay (Pierce). Proteins were reduced with 5 mM TCEP at room temperature for 15 minutes and alkylated with 15 mM Iodoacetamide at room temperature for one hour in the dark. The alkylation reaction was quenched with dithiothreitol. Proteins were precipitated using the methanol/chloroform method. In brief, four volumes of methanol, one volume of chloroform, and three volumes of water were added to the lysate, which was then vortexed and centrifuged to separate the chloroform phase from the aqueous phase. The precipitated protein was washed with one volume of ice-cold methanol. The protein pellet was allowed to air dry. Precipitated protein was resuspended in 200 mM EPPS, pH 8. Proteins were digested with LysC (1:50; enzyme:protein) overnight at 25°C followed by trypsin (1:100; enzyme:protein) for 6 hours at 37 °C. Peptide quantification was performed using the micro-BCA assay (Pierce). Equal amounts of peptide from each sample was labeled with tandem mass tag (TMT10) reagents (1:4; peptide:TMT label) (Pierce). The 10-plex labeling reactions were performed for 2 hours at 25°C. Modification of tyrosine residues with TMT was reversed by the addition of 5% hydroxyl amine for 15 minutes at 25°C. The reaction was quenched with 0.5% trifluoroacetic acid and samples were combined at a 1:1:1:1:1:1:1:1:1:1:1 ratio. Combined samples were desalted and offline fractionated into 24 fractions as previously described.

#### Liquid chromatography-MS3 spectrometry (LC-MS/MS)

12 of the 24 peptide fractions from the basic reverse phase step (every other fraction) were analyzed with an LC-MS3 data collection strategy on an Orbitrap Lumos mass spectrometer (Thermo Fisher Scientific) equipped with a Proxeon Easy nLC 1000 for online sample handling and peptide separations^124^. Approximately 5 µg of peptide resuspended in 5% formic acid + 5% acetonitrile was loaded onto a 100 µm inner diameter fused-silica micro capillary with a needle tip pulled to an internal diameter less than 5 µm. The column was packed in-house to a length of 35 cm with a C_18_ reverse phase resin (GP118 resin 1.8 μm, 120 Å, Sepax Technologies). The peptides were separated using a 180 min linear gradient from 3% to 25% buffer B (100% acetonitrile + 0.125% formic acid) equilibrated with buffer A (3% acetonitrile + 0.125% formic acid) at a flow rate of 600 nL/min across the column. The scan sequence began with an MS1 spectrum (Orbitrap analysis, resolution 120,000, 350−1350 *m/z* scan range, AGC target 1 × 10^6^, maximum injection time 100 ms, dynamic exclusion of 75 seconds). The “Top10” precursors were selected for MS2 analysis, which consisted of CID (quadrupole isolation set at 0.5 Da and ion trap analysis, AGC 1.5 × 10^4^, NCE 35, maximum injection time 150 ms). The top ten precursors from each MS2 scan were selected for MS3 analysis (synchronous precursor selection), in which precursors were fragmented by HCD prior to Orbitrap analysis (NCE 55, max AGC 1.5 × 10^5^, maximum injection time 150 ms, isolation window 2 Da, resolution 50,000).

#### LC-MS3 data analysis

A suite of in-house software tools was used for .RAW file processing and controlling peptide and protein level false discovery rates, assembling proteins from peptides, and protein quantification from peptides as previously described^124^. MS/MS spectra were searched against a Uniprot *Drosophila* reference database appended with common protein contaminants and reverse sequences. Database search criteria were as follows: tryptic with two missed cleavages, a precursor mass tolerance of 50 ppm, fragment ion mass tolerance of 1.0 Da, static alkylation of cysteine (57.02146 Da), static TMT labeling of lysine residues and N-termini of peptides (229.162932 Da), and variable oxidation of methionine (15.99491 Da). TMT reporter ion intensities were measured using a 0.003 Da window around the theoretical *m/z* for each reporter ion in the MS3 scan. Peptide spectral matches with poor quality MS3 spectra were excluded from quantitation (<200 summed signal-to-noise across 10 channels and <0.7 precursor isolation specificity).

#### Metabolomics

Three control (genotype: *elav-GAL4/+),* three tau (genotype: *elav-GAL4/+; UAS-tau^R406W^/+),* and three Aβ_1-42_ (genotype: *elav-GAL4/+; UAS-Abeta^1-42^)* samples of 40 fly heads each were collected and untargeted positively and negative charged polar and non-polar metabolites were assessed using liquid chromatography-mass spectrometry as described in detail previously^125^.

#### Identifying Age-Associated Genes in RNA-seq data from the Genotype-Tissue Expression (GTEx) project

To identify genes with significant associations between gene expression in the brain and chronological age, we sought out RNA-seq data sets with many individuals and a large dynamic range of ages. We analyzed 2642 samples from 382 individuals representing 13 different brain tissues, using the measurements of transcripts per million (TPM) available from the GTEx analysis version 8 (https://gtexportal.org/home/datasets). The age range of the patients are from 20-70 years old with a median age of 58 years old. To measure the effects of age on gene expression in the brain, we used a mixed-effects model as implemented in lme4 version 1.1.27.1, treating sex, ethnicity, patient identity and tissue of origin as covariates with the following equation:

Y ∼ Age + Sex + PMI + Tissue + Ethnicity + Sample ID

Where “Sample ID” is treated as a random effect while all other covariates are treated as fixed effects. We identify genes as significantly associated with age if the FDR-adjusted p-value for the age coefficient is less than 0.1 and if the absolute unstandardized coefficient for age is greater than 0.1, which corresponds to a change of 1 TPM per decade in this data set, assuming age is the only factor. We used this equation to assess whether there was a significant effect on gene expression with age given the mean expression of the screen hits. To assess robustness of this test, we performed 10,000 permutations of either gene sets of the same size as the set of the screen hits or over patient age. We computed an empirical p-value which was the number of permutations with p-values smaller than the original test divided by the number of permutations. When performing this analysis for individual tissues, we used a generalized additive model with the same formula but excluding the “Sample ID” and “Tissue” variables.

To perform Gene Set Enrichment Analysis, we used the R package “fgsea” version 1.14.0 using the Reactome 2022 library from Enrichr as the reference set of pathways. We added the neurodegeneration screen hits as a pathway term, for pathway enrichment analysis. We defined the gene set of this new pathway using the human orthologs of the neurodegeneration screen hits, which were mapped using the DRSC integrative ortholog prediction tool (DIOPT)^126^. We used the standardized regression coefficient to rank the genes^68,127^.

#### Analysis of single-nuclear RNA-seq data

To identify cellular subtypes that were associated with disease, we analyzed previously published single nuclear RNA-seq data^66^, which included 70,000 cells from 24 Alzheimer’s disease patients and 24 age and sex-matched healthy controls. The data were preprocessed as in previous work^66^. In short, for each of the previously defined “broad cell types” (excitatory neurons, inhibitory neurons, astrocytes, oligodendrocytes, microglia and oligodendrocyte progenitor cells), we applied Seurat version 4.0.4’s implementations for log-normalizing the data, detecting highly variable features, and standard scaling the data. We used Seurat’s implementation of PCA reducing the data to 20 principal components. After applying PCA, we used Harmony version 0.1 to correct for the effects of sex, individual, sequencing batch and post-mortem interval in our data. This correction was performed to minimize the effects of confounders in our clustering analysis. We further applied Scrublet to predict and remove doublet cells from the population as implemented in Scanpy version 1.8.2. We used the Harmony components for UMAP dimensionality reduction and Leiden clustering. To determine the Leiden clustering resolution, we calculated the silhouette coefficient after applying Leiden clustering on a range of values (resolution={0.1,0.2,0.3,0.4, 0.6, 0.8, 1.0, 1.4, 1.6, 2.0,2.1,2.2,2.3,2.4,2.5}). We selected the clustering resolution that maximized the silhouette coefficient. To identify disease-associated clusters, we applied a hypergeometric test to determine if a cluster was over-represented by cells derived from Alzheimer’s disease patients or healthy controls. We subsequently applied MAST as implemented in Seurat to determine the differentially expressed genes between Alzheimer’s disease-enriched clusters and the remaining sub-clusters within a given cell type. We defined differentially expressed genes as having an FDR-adjusted p-value less than 0.1 and an absolute log_2_ fold change greater than 1.

#### *Analysis* of Drosophila *multi-omics*

We performed two-way t-tests to assess the significance of *Drosophila* proteins, phosphoproteins and metabolites between *Drosophila* models of amyloid β and control as well as significant proteins, phosphoproteins and metabolites between *Drosophila* models of tau and control. We used gProfiler with the g_SCS multiple hypothesis correction to identify significant gene ontology terms using *Drosophila* pathways as a reference^128^. We used PiuMet to map unannotated m/z peaks in the metabolomic data to known compounds^129^.

#### Laser-capture RNA-seq

We used the laser-capture RNA-seq method to profile total RNA of brain neurons similar to what we reported in previous studies^37,38^. In brief, laser-capture microdissection was performed on human autopsy brain samples to extract neurons^38^. For each temporal cortex (middle gyrus) about 300 pyramidal neurons were outlined in layers V/VI by their characteristic size, shape, and location in HistoGene-stained frozen sections and laser-captured using the Arcturus Veritas Microdissection System (Applied Biosystems) as in previous studies^38^. Linear amplification, construction, quantification, and quality control of sequencing libraries, fragmentation, and sequencing methods were described in earlier studies^38^. RNA seq data processing and quality control was performed similar to what we reported in previous studies^37,38^. In summary,

The RNA-sequencing data was aligned to the human genome reference sequence hg19 using TopHat v2.0 and Cufflinks v1.3.0. To measure RNA-sequencing quality control, we used FASTQC and RNA-SeQC. We blinded ourselves to the disease status of the patient when preparing the samples.

#### Data sets used for expression Quantitative Trait Locus (eQTL) analysis

eQTL analysis was performed using seven previously published bulk cortex data sets and one new laser-captured pyramidal neuron data set. ROSMAP, MayoRNAseq, MSBB, and HBTRC data were obtained from the AD Knowledge Portal (https://adknowledgeportal.org) on the Synapse platform (Synapse ID: syn9702085). CommonMind was obtained from the CommonMind Consortium Knowledge Portal (https://doi.org/10.7303/syn2759792) also on the Synapse platform (Synapse ID: syn2759792); GTEx was obtained from https://gtexportal.org/home/. UKBEC, was obtained from http://www.braineac.org/; BRAINCODE, was obtained from http://www.humanbraincode.org/. The data sets are described in detail at each of the source portals and in the corresponding original publications^37,38,130–138^.

We used a conservative four-stage design: ***1***, Cortex discovery stage: eQTL analysis in four human cortex cohorts (stage D in Supplementary Table 1). ***2***, Cortex replication stage: replication of findings from the discovery stage in three independent human cortex cohorts (stage R in Supplementary Table 1). ***3***, To further enhance statistical power, we performed meta-analysis across all seven cohorts. This meta-analysis highlighted an additional 17 suggestive eGenes with *P* values < 5 * 10^-8^ (Table S2) which were not recovered in the two-stage design. 4, We confirmed 12 suggestive eGenes in the laser-captured pyramidal neuron data set with P values < 0.05.

#### Gene expression data processing

For RNAseq data sets, the gene reads counts were normalized to TPM (Transcripts Per Kilobase Million) by scaling gene length first and followed by scaling sequencing depth. The gene length was considered as the union of exon lengths. Consistent and stringent quality control and normalization steps were applied for each of the cohorts: 1) For sample quality control, we removed samples with poor alignment. We kept samples with > 10M mapped reads and > 70% mappability by considering reads with mapping quality of Q20 or higher (the estimated read alignment error rate was 0.01 or less). 2) Filtering sample mix-ups by comparing the reported sex with the transcriptional sex determined by the expression of female-specific *XIST* gene and male-specific *RPS4Y1* gene. 3) Filtering sample outliers. Sample outliers with problematic gene expression profiles were detected based on Relative Log Expression (RLE) analysis, spearman correlation based hierarchical clustering, D-statistics analysis^139^. 4) For normalization, gene expression values were quantile normalized after log10 transformed by adding a pseudo count of 1e-4. 5) SVA package was applied for removing batch effects by using combat function and adjusting age, sex, RIN, PMI. We accounted for latent covariates by performing fsva function. Residuals were outputted for downstream analysis. For array-based gene expression datasets, we directly used the downloaded, quality-controlled gene expression profiles.

#### Genotype data processing for eQTL analyses

We applied PLINK2 (v1.9beta) and in house scripts to perform rigorous subject and SNP quality control (QC) for each dataset in the following order: 1) Set heterozygous haploid genotypes as missing; 2) remove subjects with call rate < 95%; 3) remove subjects with gender misidentification; 4) remove SNPs with genotype call rate < 95%; 5) remove SNPs with Hardy-Weinberg Equilibrium testing P value < 1 * 10^-6^; 6) remove SNPs with informative missingness test (Test-mishap) P value < 1 * 10^-9^; 7) remove SNPs with minor allele frequency (MAF) < 0.05; 8) remove subjects with outlying heterozygosity rate based on heterozygosity F score (beyond 4*sd from the mean F score); 9) IBS/IBD filtering: pairwise identity-by-state probabilities were computed for removing both individuals in each pair with IBD>0.98 and one subject of each pair with IBD > 0.1875; (10) population substructure was tested by performing principal component analysis (PCA) using smartPCA in EIGENSOFT^140^. PCA outliers were excluded and the top 3 principal components were used as covariates for adjusting population substructures.

#### Imputation of Genotypes for eQTL analyses

The array-based genotype datasets were enhanced by genotype imputation. Genotype imputation for each dataset was performed separately on Michigan Imputation Server, using 1000G phase 3 reference panel. Eagle v2.3 and Minimac3 were selected for phasing and imputing respectively, and EUR population was selected for QC. Only variants with R^2^ estimates less than or equal to 0.3 were excluded from further analysis. And only variants with MAF > 5% were also included in downstream eQTL analysis. Prior to imputation, pre-imputation checks provided by Will Rayner performed external quality controls to fit the requirements of the imputation server.

We used European population reference (EUR) haplotype data from the 1000 Genomes Project (2010 interim release based on sequence data freeze from 4 August 2010 and phased haplotypes from December 2010) to impute genotypes for up to 6,709,258 SNPs per data set. We excluded SNPs that did not pass quality control in each study

#### eQTL analysis

The eQTL mapping was conducted using R Package Matrix EQTL using the additive linear model on a high-performance Linux-based Orchestra cluster at Harvard Medical School. For cis-eQTL analysis, SNPs were included if their positions were within 1Mb with the TSS of a gene. And trans-eQTL analysis included SNP-gene association if their distance was beyond this window. FDRs reported by MatrixEQTL were used to estimate the association between SNPs and gene expression.

#### Meta eQTL analysis

We performed meta eQTL analysis using three separate effects model implemented in METASOFT^141^, which took effect size and standard error of SNP-gene pair in each dataset as input. Fixed effects model (FE model) was based on inverse-variance-weighted effect sizes. Random Effects model (RE model) was a very conservative model based on inverse-variance-weighted effect size. Han and Eskin’s random effects model (RE2 model) was optimized to detect associations under heterogeneity. We reported statistics of the RE2 model in this study.

#### Identifying eGene-associated variants that associate with transcription factor binding

We were interested in determining whether eGene-associated variants overlapped with transcription factor binding sites. We used the optimal hg19 ChIP-seq-derived transcription factor peak sets from ENCODE 3, which we downloaded from the UCSC genome browser. To determine if the eQTL of interest overlapped with a DNA-binding motif, we extracted the sequence 50 base pairs upstream and 50 base pairs downstream of the variant and used FIMO to detect the presence of an overlapping DNA-binding motif^142^. We used the HOCOMOCO version 11 core motif set as reference motifs. Correlations between *HLA-DRB1* and *CUX1* expression were performed using Pearson’s correlation test as implemented in R version 4.0.2.

To identify correlations between eGenes and biological pathways, we applied GSVA version 1.42.0 to the CPM-normalized temporal cortex pyramidal neuron RNA-seq to identify the REACTOME pathway enrichments per-sample. For this analysis we used the REACTOME pathways available in GSVAdata version 1.30, which downloads the REACTOME pathways from msigdb version 3.0 with the data set named “c2BroadSets”. We calculated correlations between GSVA signatures and gene expression using the Pearson correlation coefficient as implemented in R version 4.0.2, considering correlations with an FDR-adjusted p-value less than 0.1.

#### Design of integration analyses

In order to identify the biological mechanisms through which human and model organism genetic hits contribute to neurodegenerative disease, we utilized the Prize-Collecting Steiner Forest algorithm (PCSF) as implemented in OmicsIntegrator 2^42^ (OI v2.3.10, https://github.com/fraenkel-lab/OmicsIntegrator2). The PCSF algorithm identifies disease-associated networks based on signals derived from molecular data that are significantly altered in individuals with the disease. The algorithm aims to retrieve as many termini as possible while minimizing the number of low-confidence edges in the final network. The resulting network includes a subset of the input nodes and high-confidence interactors that were not part of the input. We used OmicsIntegrator to map proteomic, phosphoproteomic, metabolomic and genetic changes to a set of known protein-protein and protein-metabolite interactions derived from physical protein-protein interactions from iRefIndex version 17 and protein-metabolite interactions described in the HMDB and Recon 2 databases. To add brain-specific edges, we include the interactions derived from Affinity Purification Mass Spectrometry (AP-MS) of mice in BraInMap^143^. Additionally, we include previously published interactions between proteins found in tau aggregates and phosphorylated tau derived from AP-MS of neurofibrillary tangles^144^. The costs on the protein-protein interactions were computed as 1 minus the edge score reported by iRefIndex, while the cost of the protein-metabolite interactions were calculated as in previous studies^129,145^. Given that these reference interactions were defined in human proteins and metabolites, we mapped the *Drosophila* proteins and phosphoproteins to their human orthologs using DIOPT version 8.0, choosing the human orthologs that the tool determined were of “moderate” or “high” confidence^126^ (https://www.flyrnai.org/cgi-bin/DRSC_orthologs.pl). Briefly, DIOPT identifies gene-ortholog pairs from multiple databases and computes a score based on how many databases report the gene-ortholog pair. We chose the human ortholog most frequently reported by this ensemble of databases. Orthologs reported in fewer than three reference databases were also excluded. In order to comprehensively characterize metabolomic changes, we used PiuMet to map uncharacterized metabolites to compounds identified in HMDB^129^. In addition to integrating the phosphoproteomic data, we included the predicted upstream kinases from iProteinDB whose proteomic levels in *Drosophila* correlated with its phosphoproteomic targets^146^ (Spearman correlation coefficient>0.4, https://www.flyrnai.org/tools/iproteindb/web/).

For the Alzheimer’s disease-specific network, we integrated the screen hits with genetic modifiers of disease severity from model organism screens and available proteomics, phosphoproteomics and metabolomics from the literature and generated data. These data are derived from multiple brain tissues, and several brain regions are represented in individual data sets as well (Supplementary Table 7). To make sure no individual brain region or study was overrepresented in terms of the number of weighted nodes, we applied different thresholds of significance for each data source to generate the Alzheimer’s disease network. The prizes of the proteomic, phosphoproteomic, and metabolomic data are calculated as the negative base 10 logarithm of the Benjamini-Hochberg FDR-corrected p-value calculated by a two-way t-test. The *Drosophila* phosphoproteomic data and the metabolomic data were subject to an FDR threshold of 0.1, while the *Drosophila* proteomic and human proteomic data had more stringent cutoffs (FDR<0.01 and FDR<0.0001 respectively). Additionally, the metabolomic data were only assigned prizes if the absolute log_2_ fold change was greater than 1. The human lipidomic data were assigned prizes by their negative log_10_ nominal p-value and were included if their nominal p-value was less than 0.05. The upstream kinases were assigned the same prizes as the targeted phosphoproteins. Instead of assigning prizes based on a two-way t-test, the genetic hits were assigned prizes differently. For the human eGenes, prizes were assigned to all genes in the discovery phase with a value equal to -log_10_(genome-wide FDR). For genes found in the *Drosophila* neurodegeneration and tau aggregation screens, prizes were set to 1=-log_10_(0.1). For the human GWAS loci, the prizes were set to the -log_10_ Bonferroni corrected, genome-wide p-value for those determined to be causal loci according to previous analyses and 1 otherwise^9^. After the initial prize assignments, the values are minimum-maximum normalized to values between 0 and 1 within each data type, weighing each prize by a scale factor reflecting our confidence in the degree to which a given data type reflects Alzheimer’s disease pathology (Supplementary Table 7). We further included previously published *Drosophila* modifiers of tau toxicity^15^. If any nodes overlapped between data sets, we attributed the node to the data set for which it was assigned the greatest weight and dropped all entries with smaller weights for the given node.

For each network, we performed 100 randomizations of the edges with gaussian noise to assess the robustness of the nodes to perturbations to edges and prize values. Additionally, we performed 100 randomizations of the prize values to assess the specificity of each node to their assigned prizes. We filtered out nodes that did not have a prize (Steiner nodes) if they appeared in fewer than 40 of the robustness randomizations and more than 40 of the specificity randomizations. We then performed Louvain clustering with a resolution of 1.0 for community detection in the networks. We applied Leiden clustering with the same resolution parameter and calculated the modularity scores for Louvain and Leiden clustering results. Louvain clustering had a slightly higher Q-value than Leiden clustering, so we reported the Louvain clustering results for our network analysis (Louvain Q=0.415, Leiden Q=0.412).

OmicsIntegrator hyperparameters control the weights on prizes (β), the weight of the edges on the dummy node for network size (ω) and the edge penalty for highly connected “hub” nodes (γ). In order to select hyperparameters for OmicsIntegrator, we evaluated a range of hyperparameters for each network: β={2,5,10}, ω={1,3,6} and γ={2,5,6}. We chose networks based on minimizing the mean specificity, maximizing the mean robustness and minimizing the KS statistic between node degree of the prizes as compared to those of the predicted nodes. Our final selection was β=5, ω=3 and γ=5.

Networks were visualized using Cytoscape version 3.8.0.

#### Implementation of ROBUST to compare to the result of OmicsIntegrator

We applied ROBUST to the same interactome and initial node list as we did for OmicsIntegrator. We used the version of ROBUST from https://github.com/bionetslab/robust_bias_aware/tree/main. We used the default parameters for ROBUST.

#### Comparison of OmicsIntegrator to a graph neural network and GNNExplainer

We trained a graph neural network with two graph convolutional layers using a mean squared error loss function to predict the node weights supplied to OmicsIntegrator using the same interactome as we used for OmicsIntegrator. The graph neural network was implemented in Pytorch Geometric version 2.5.2. To select hyperparameters, we trained the algorithm on 80% of the nodes and tested the model on 20% of the data selecting the combination of learning rate, weight decay and number of epochs minimized the loss between the predicted and observed values. We found that the best learning rate was 0.01, the optimal number of epochs was 100 and the best weight decay was 1*10^-4^. We re-ran the model on our whole data set using these hyperparameters and applied GNNExplainer as implemented in Pytorch Geometric version 2.5.2. To establish which node attributions were most important for prediction, we re-ran the model after re-distributing node weights to establish a null distribution of node attributions. We kept the nodes that were within the top 99^th^ quantile of the empirical distribution.

#### Semiquantitative scale for assaying the Drosophila rough eye phenotype

The approach for evaluating the *Drosophila* rough eye phenotype is described in^24^. We used a semiquantitative scale ranging from 0-5. 0 represents a wild-type eye, 1 refers to less than 50% of fly eye ommatidia displaying morphological disruption. A score of 2 indicates that there is morphological disruption in 50-100% of the fly eye ommatidia and 0-25% reduction in eye size. A score of 3 indicates 100% ommatidial disruption and a 25-50% in fly eye size. A score of 4 describes fly eyes with 100% ommatidial disruption and one of the following features: ommatidial fusions, darkened or discolored areas, or over 50% reduction in eye size. A score of 5 refers to eyes with at least two of the features that can be observed in an eye with a rough eye score of 4.

#### Betweenness analysis of neurodegeneration screen hits with regulators of DNA damage

We used the R package igraph version 1.2.6 to compute the subnetwork of neurodegeneration screen hits, and regulators of DNA damage. We used the package’s implementation of betweenness centrality to compute each node’s betweenness in this subnetwork.

#### COMET assay for DNA damage in human neural progenitor cells

For the alkaline COMET assay, we applied inhibitors of CK2 (CX-4945) and the NOTCH signaling pathway (Compound E) to human iPSC-derived neural progenitor cells. Cells were treated with 5µM of the inhibitor overnight and harvested. Comets were selected using the OpenComet plugin in ImageJ^111,147^. The extent of DNA damage was measured by the tail moment and proportion of intensity between the tail and the head of the comet. The tail represents single-stranded DNA that trails off from the nucleus due to DNA damage burden. Longer tails indicate a greater extent of DNA damage. DMSO and etoposide were included as negative and positive controls respectively.

#### Human iPSC culture

Human iPSCs (male WTC11 background, gift from the lab of Michael Ward) harboring a single-copy of doxycycline-inducible (DOX) mouse NGN2 at the AAVS1 locus and pC13N-dCas9-BFP-KRAB at human CLYBL intragenic safe harbor locus (between exons 2 and 3) were cultured in mTeSR Medium (Stemcell Technologies; Cat. No. 85850) on Corning Tissue Culture Treated Dishes (VWR International LLC; Cat. No. 62406-161) coated with Geltrex (Life Technologies Corporation; Cat. No. A1413301). Briefly, mTESR medium supplemented with mTESR supplement (Stemcell Technologies; Cat. No. 85850) and antibiotic Normocin (Invivogen; Cat. No. Ant-nr-2) was replenished every day until 50% confluent^148^. Stem cells were transduced with lentivirus packaged with CROPseq-Guide-Puro vectors and selected with Puromycin (Gibco; Cat. No. A11138-03) until confluent. When cells were 80%-90% confluent, cells were passaged, which entailed the following: dissociating in Accutase (Stemcell Technologies; Cat. No. 7920) at 37°C for 5 minutes, diluting Accutase 1:5 with mTeSR medium, collecting in conicals and centrifuging at 300g for 5 minutes, asipirating supernatant, resuspending in mTESR supplemented with 10uM Y-27632 dihydrochloride ROCK inhibitor (Stemcell Technologies; Cat. No. 72302) and plating in Geltrex-coated plates.

#### NGN2 Neuronal Differentiation and RNA extraction

Neuronal differentiation was performed as described in previous work with a few modifications^149^. On day 1, iPSCs transduced with CROPseq-Guide-Puro were plated at 40,000 cells/cm^2^ density in Geltrex-coated tissue culture plates in mTESR medium supplemented with ROCK inhibitor and 2µg/ml Doxycycline hyclate (Sigma; Cat. No. D9891-25G). On Day 2, Medium was replaced with Neuronal Induction media containing the following: DMEM/F12 (Gibco; Cat. No 11320-033), N2 supplement (Life Technologies Corporation; Cat. No. 17502048), 20% Dextrose (VWR; Cat. No. BDH9230-500G), Glutamax (Life Technologies Corporation; Cat. No. 35050079), Normocin (Invivogen; Cat. No. Ant-nr-2), 100 nM LDN-193189 (Stemcell Technologies; Cat. No. 72147), 10uM SB431542 (Stemcell Technologies; Cat. No. 72234) and 2uM XAV (Stemcell Technologies; Cat. No. 72674) and 2µg/ml DOX. The Neuronal Induction Media was replenished on day 3. On day 4, the medium was replaced with Neurobasal Media (Life Technologies Corporation; Cat. No. 21103049) containing B27 supplement (Gibco; Cat. No. 17504044), MEM Non-Essential Amino Acids (Life Technologies Corporation; Cat. No. 11140076) Glutamax, 20% Dextrose, 2µg/ml DOX, Normocin, 10ng/ml BDNF (R&D Systems; Cat. No. 11166-BD), 10ng/ml CNTF (R&D Systems; Cat. No. 257-NT/CF), 10ng/ml GDNF (R&D Systems; Cat. No. 212-GD/CF) and cultured for 2 days. At day 6, cells were dissociated with Accutase and resuspended with Trizol (Thermofisher Scientific; Cat. No.15596018). RNA was extracted following manufacturer’s manual, using Direct-zol RNA Miniprep kit (Zymo Research, R2050)

#### Bulk RNA-seq analysis of CRISPRi knockdowns in neural progenitor cells

We analyzed the RNA-seq data after CRISPRi knockdown as performed in previous CRISPRi studies^150^. In summary, we mapped the raw sequencing reads to the hg38 reference transcriptome with salmon version 1.10.1. We used the ‘-noLengthCorrection’ option to generate transcript abundance counts. We generated gene-level count estimates with tximport version 1.16.1. To account for the effects of different guides, we performed differential expression analysis between knockdown and control with DESeq2 version 1.28.1 treating guide identity as a covariate. We applied the apelgm package version 1.10.0 to shrink the log_2_ fold changes. We applied Gene Set Enrichment Analysis to the ranked, shrunk log_2_ fold changes using the fgsea package version 1.14.0 and the Reactome 2022 library as the reference pathway set^68,127^.

## Acknowledgements

We thank Leslie Gaffney for assistance with figure design and thank all members of the Fraenkel lab for their feedback on the project and manuscript. We thank the MIT BioMicroCenter for RNA sequencing of the NGN2 neural progenitor cells. M.J.L. was supported in part by the Barbara Weedon Fellowship. Fly stocks were obtained from the Bloomington Drosophila Stock Center (NIH P40-OD018537) and the Vienna Drosophila Resource Center. We thank the Transgenic RNAi Project (TRiP) at Harvard Medical School (NIH R24 OD030002; PI: N. Perrimon) for making Drosophila stocks^151^. Monoclonal antibodies were obtained from the Developmental Studies Hybridoma Bank developed under the auspices of the NICHD and maintained by the University of Iowa, Department of Biology, Iowa City, IA 52242. Yi Zhong provided excellent technical assistance. This research was funded by NIH R01-AG057331 to E.F., C.R.S., and M.B.F. The results published here are in whole or in part based on data obtained from the AD Knowledge Portal (https://adknowledgeportal.org). The ROSMAP study data were provided by the Rush Alzheimer’s Disease Center, Rush University Medical Center, Chicago. Data collection was supported through funding by NIA grants P30AG10161 (ROS), R01AG15819 (ROSMAP; genomics and RNAseq), R01AG17917 (MAP), R01AG36836 (RNAseq), U01AG32984 (genomic and whole exome sequencing), U01AG61356 (whole genome sequencing, targeted proteomics, ROSMAP AMP-AD), the Illinois Department of Public Health (ROSMAP), and the Translational Genomics Research Institute (genomic). The MSBB data were generated from postmortem brain tissue collected through the Mount Sinai VA Medical Center Brain Bank and were provided by Dr. Eric Schadt from Mount Sinai School of Medicine. The Mayo RNAseq study data was led by Dr. Nilüfer Ertekin-Taner, Mayo Clinic, Jacksonville, FL as part of the multi-PI U01 AG046139 (MPIs Golde, Ertekin-Taner, Younkin, Price). Samples were provided from the following sources: The Mayo Clinic Brain Bank. Data collection was supported through funding by NIA grants P50 AG016574, R01 AG032990, U01 AG046139, R01 AG018023, U01 AG006576, U01 AG006786, R01 AG025711, R01 AG017216, R01 AG003949, NINDS grant R01 NS080820, CurePSP Foundation, and support from Mayo Foundation. Study data includes samples collected through the Sun Health Research Institute Brain and Body Donation Program of Sun City, Arizona. The Brain and Body Donation Program is supported by the National Institute of Neurological Disorders and Stroke (U24 NS072026 National Brain and Tissue Resource for Parkinsons Disease and Related Disorders), the National Institute on Aging (P30 AG19610 Arizona Alzheimers Disease Core Center), the Arizona Department of Health Services (contract 211002, Arizona Alzheimers Research Center), the Arizona Biomedical Research Commission (contracts 4001, 0011, 05-901 and 1001 to the Arizona Parkinson’s Disease Consortium) and the Michael J. Fox Foundation for Parkinsons Research. For the CommonMind study, Data were generated as part of the CommonMind Consortium supported by funding from Takeda Pharmaceuticals Company Limited, F. Hoffman-La Roche Ltd and NIH grants R01MH085542, R01MH093725, P50MH066392, P50MH080405, R01MH097276, RO1-MH-075916, P50M096891, P50MH084053S1, R37MH057881, AG02219, AG05138, MH06692, R01MH110921, R01MH109677, R01MH109897, U01MH103392, U01MH116442, project ZIC MH002903 and contract HHSN271201300031C through IRP NIMH. Brain tissue for the study was obtained from the following brain bank collections: The Mount Sinai/JJ Peters VA Medical Center NIH Brain and Tissue Repository, the University of Pennsylvania Alzheimer’s Disease Core Center, the University of Pittsburgh Brain Tissue Donation Program, and the NIMH Human Brain Collection Core. CMC Leadership: Panos Roussos, Joseph Buxbaum, Andrew Chess, Schahram Akbarian, Vahram Haroutunian (Icahn School of Medicine at Mount Sinai), Bernie Devlin, David Lewis (University of Pittsburgh), Raquel Gur (University of Pennsylvania), Chang-Gyu Hahn (Thomas Jefferson University), Enrico Domenici (University of Trento), Mette A. Peters, Solveig Sieberts (Sage Bionetworks), Stefano Marenco, Barbara K. Lipska, Francis J. McMahon (NIMH).

## Author contributions

M.J.L., C.R.S., M.B.F., and E.F. conceived and designed the study. C.A.Z, H.B., and M.B.F, designed and performed all experiments in *Drosophila*. M.J.L. performed computational analysis of published human bulk and single-cell RNA-seq, differential *Drosophila* proteomics, phosphoproteomics and metabolomics, multi-omic network integration, and RNA-seq analysis of iPSC-derived neural progenitor cells. J.P.,T.W.,D.G.,Z.L., X.D., and C.R.S. designed and performed laser-capture microdissection and eQTL meta-analysis. P-C.P. and L-H.T. designed and performed COMET assay in iPSC-derived neural progenitor cells. B.K., S.D., J.B., R.N., and S.F. designed and performed iPSC-derived NGN2 neuron differentiation and RNA-seq experiments. M.J.L., M.B.F., and E.F. wrote the manuscript. C.R.S., M.B.F., and E.F. provided funding for the study.

## Ethics Declarations

### Ethical acquisition of human patient data

Human patient data was acquired through data use agreements with the AD Knowledge Portal and the Genotype-Tissue Expression Project. We complied with the data use agreements that ensured the data were collected with informed consent from the patients in the studies. Human patient brains for laser capture microdissection were recruited with the oversight of the Institutional Review Board of Brigham and Women’s hospital.

## Supplementary Information

**Supplementary Table 1, related to Fig. 2.** GO biological process pathway enrichment analysis results for *Drosophila* modifiers of age-associated neurodegeneration. The second sheet shows the GO biological pathway enrichment analysis for all tested *Drosophila* genes. The third sheet lists all *Drosophila* stocks tested.

**Supplementary Table 2, related to Fig. 3.** eQTL analysis was performed in eight cohorts based on cortex and laser captured temporal cortex pyramidal neurons (TCPY). Abbreviations: D, Discovery phase; TCPY, temporal cortex pyramidal neurons; R, Replication phase; RIN, RNA Integrity Number; PMI, Post-mortem interval; sd, standard deviation; WGS, Whole Genome Sequencing.

**Supplementary Table 3, related to Table 1**: Results from the discovery phase of the eQTL analysis.

**Supplementary Table 4, related to Fig. 3:** Proteomics from control flies, Aβ_1-42_ transgenic flies, and tau^R406W^ transgenic flies.

**Supplementary Table 5, related to Fig. 3:** Phosphoproteomics from control flies, Aβ_1-42_ transgenic flies, and tau^R406W^ transgenic flies.

**Supplementary Table 6, related to Fig. 3:** Metabolomics from control flies, Aβ_1-42_ transgenic flies, and tau^R406W^ transgenic flies.

**Supplementary Table 7, related to Fig. 4:** Files regarding network inputs and outputs. The first sheet is the Input file for OmicsIntegrator2 analysis. “ID” indicates the gene or metabolite name, “prize_val” is the min-max normalized weight for the individual node, “source” is the data type of origin and “magnitude” is the effect size of change, where available. “Node prize overlap” shows the nodes in the network enriched in multiple data sources, the data sources from which they were derived and the normalized prize values per group. “Nodes in network” shows the nodes in the final network solution. “Edges contributed” shows the number of edges that include a particular data source. Pathways by subnetwork shows the enriched pathways in each subnetwork. “Contributed pathways” shows the pathways that were contributed by a particular data type, while contributed nodes shows the pathways that were contributed by a particular data type. “Node count by parameter” shows the number of nodes in the final network solutions depending on the hyperparameter choices. “Pathway enrichments overlapping” shows the results of hypergeometric tests for pathway enrichment in nodes that are enriched in more than one data type. “Pathway enrichments by method” shows the pathway enrichments using different network optimization methods.

**Supplementary Table 8, related to Figs. 5 and 6:** Results from the differential expression analyses for *NOTCH1*, *CSNK2A1* and *HNRNPA2B1* knockdown in NGN2 neural progenitor cells.

**Supplementary Table 9, related to Figs. 5 and 6:** Gene Set Enrichment analysis results for NGN2 neural progenitor cells after *NOTCH1*, *CSNK2A1* and *HNRNPA2B1* knockdown.

**Supplementary Table 10, related to Fig. 6:** Measurement of betweenness centrality among nodes that regulate DNA damage and neurodegeneration screen hits.

**Supplementary Fig. 1:**
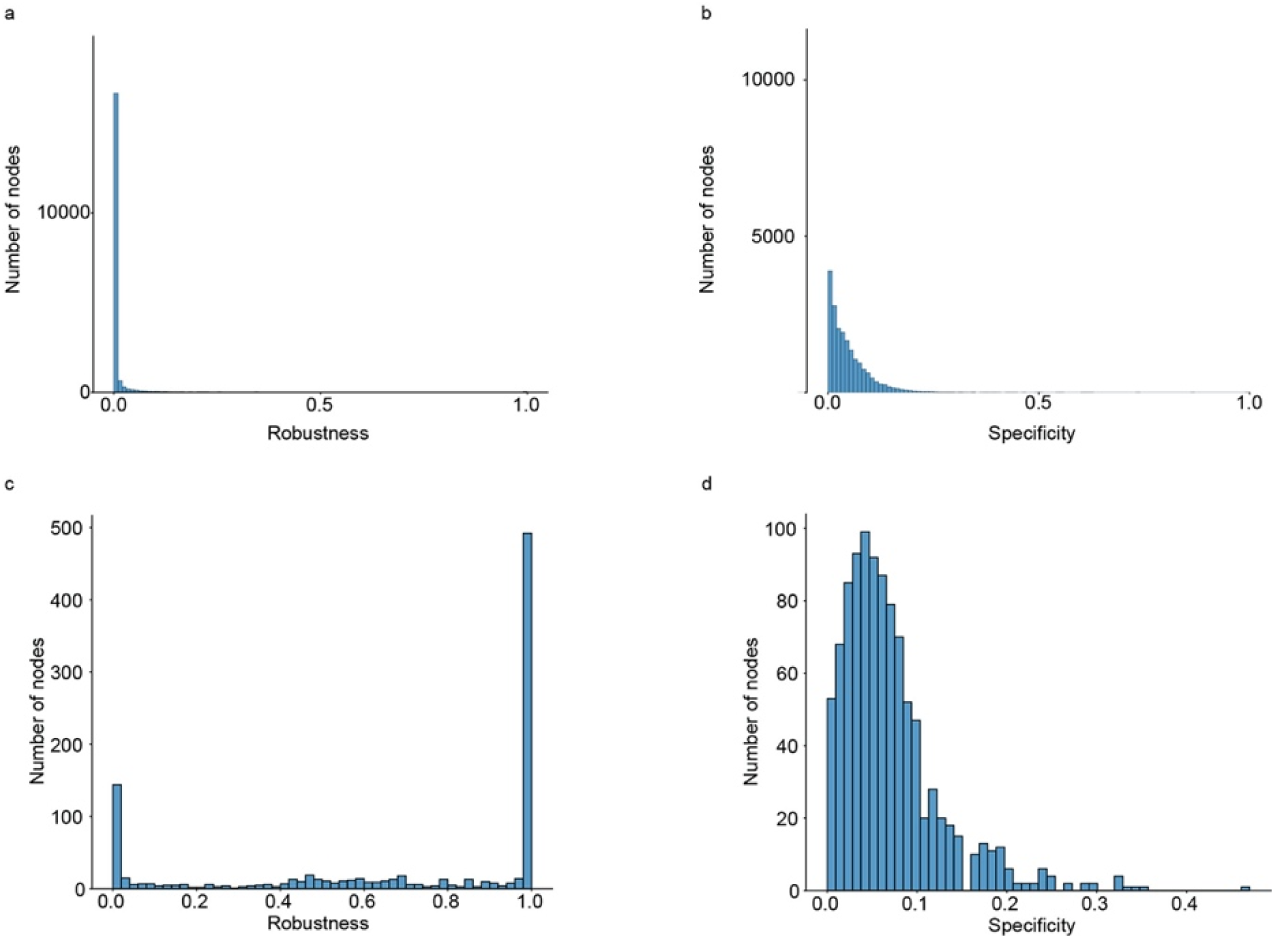
Histograms showing the distribution of a) node robustness and b) node specificity after randomizations. 0 indicates low robustness and 1 indicates high robustness, while 0 indicates high specificity and 1 indicates low specificity. After removing nodes for a robustness of at least 0.4 and a specificity of at most 0.4, we filtered out 19102 nodes. c) node robustness and d) node specificity distributions of nodes after filtering predicted nodes for insufficient robustness or specificity.

## Extended Data

**Extended Data Fig. 1, related to Fig. 2:**
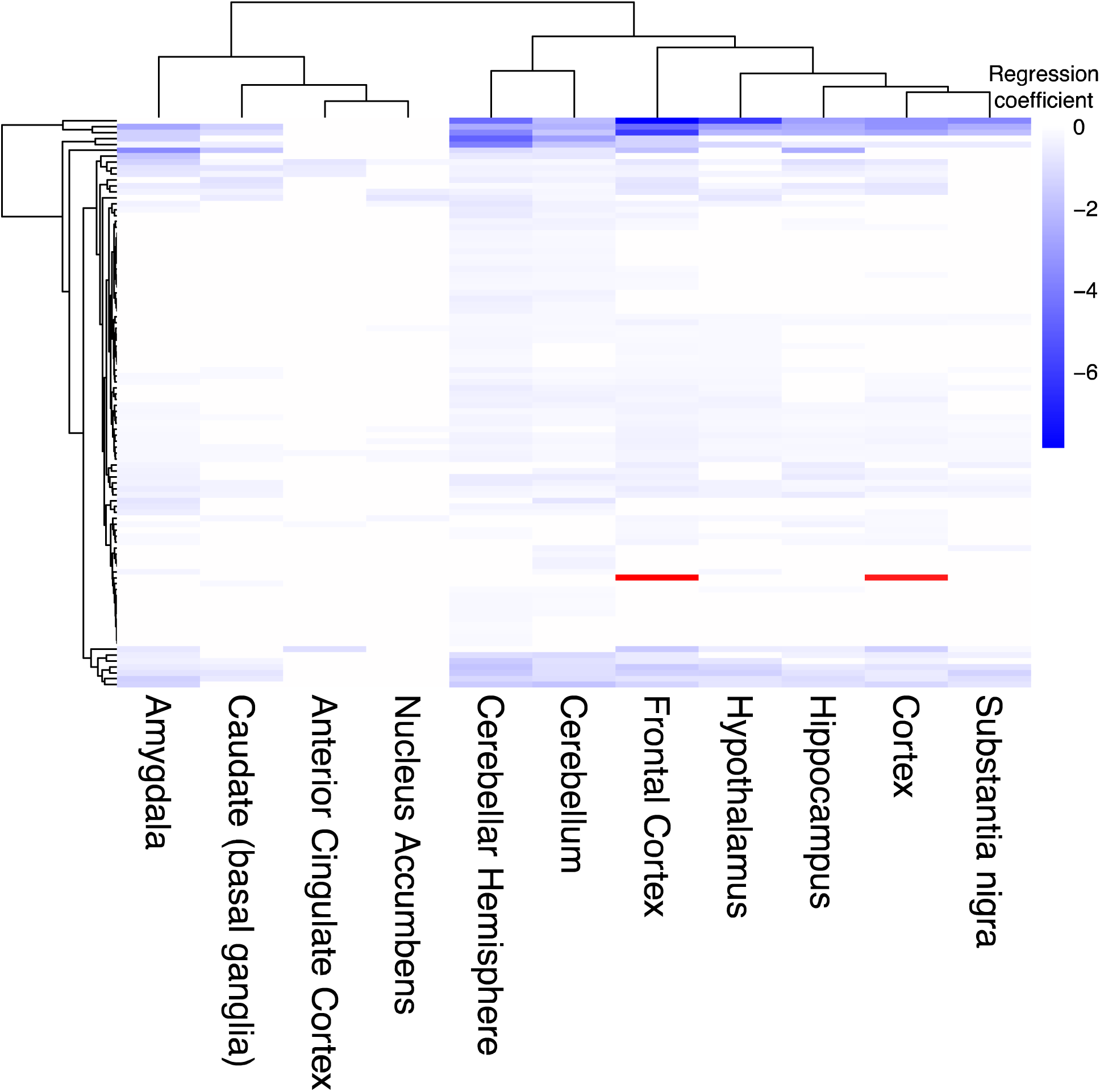
Neurodegeneration screen hits have significant changes in gene expression with respect to age across multiple brain tissues. Heatmap depicts significant linear mixed model regression coefficients between the expression of neurodegeneration screen hits and patient age in human RNA-seq in the Genotype-Tissue Expression project (GTEx) for each brain tissue. Each row is an age-associated neurodegeneration gene while each column indicates the brain tissue in GTEx, grouped by hierarchical clustering. Blue indicates a negative association and red indicates a positive association between gene expression in age as measured by the model regression coefficient. The gene with a positive regression coefficient is *HES6*.

**Extended Data Fig. 2, related to Fig. 2:**
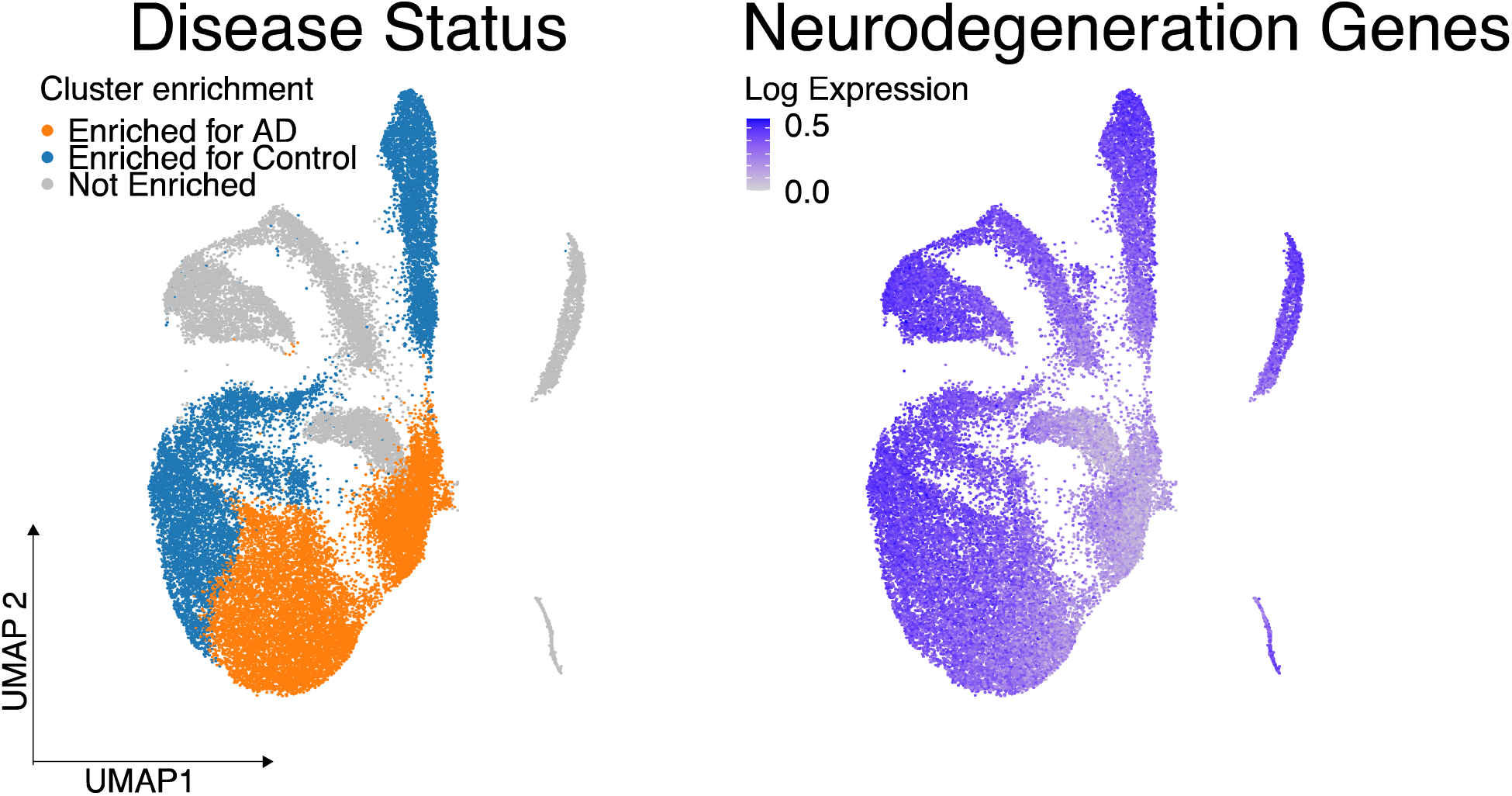
The average expression of age-associated neurodegeneration genes declines in Alzheimer’s disease-associated excitatory neurons. UMAP projections depict excitatory neurons from Mathys et al. 2019. In the left plot, cells are shaded by whether they belong to clusters overrepresented by cells from control or Alzheimer’s disease patients. The right UMAP shows the average expression of age-associated neurodegeneration genes in this group of excitatory neurons.

**Extended Data Fig. 3, related to Fig. 3.**
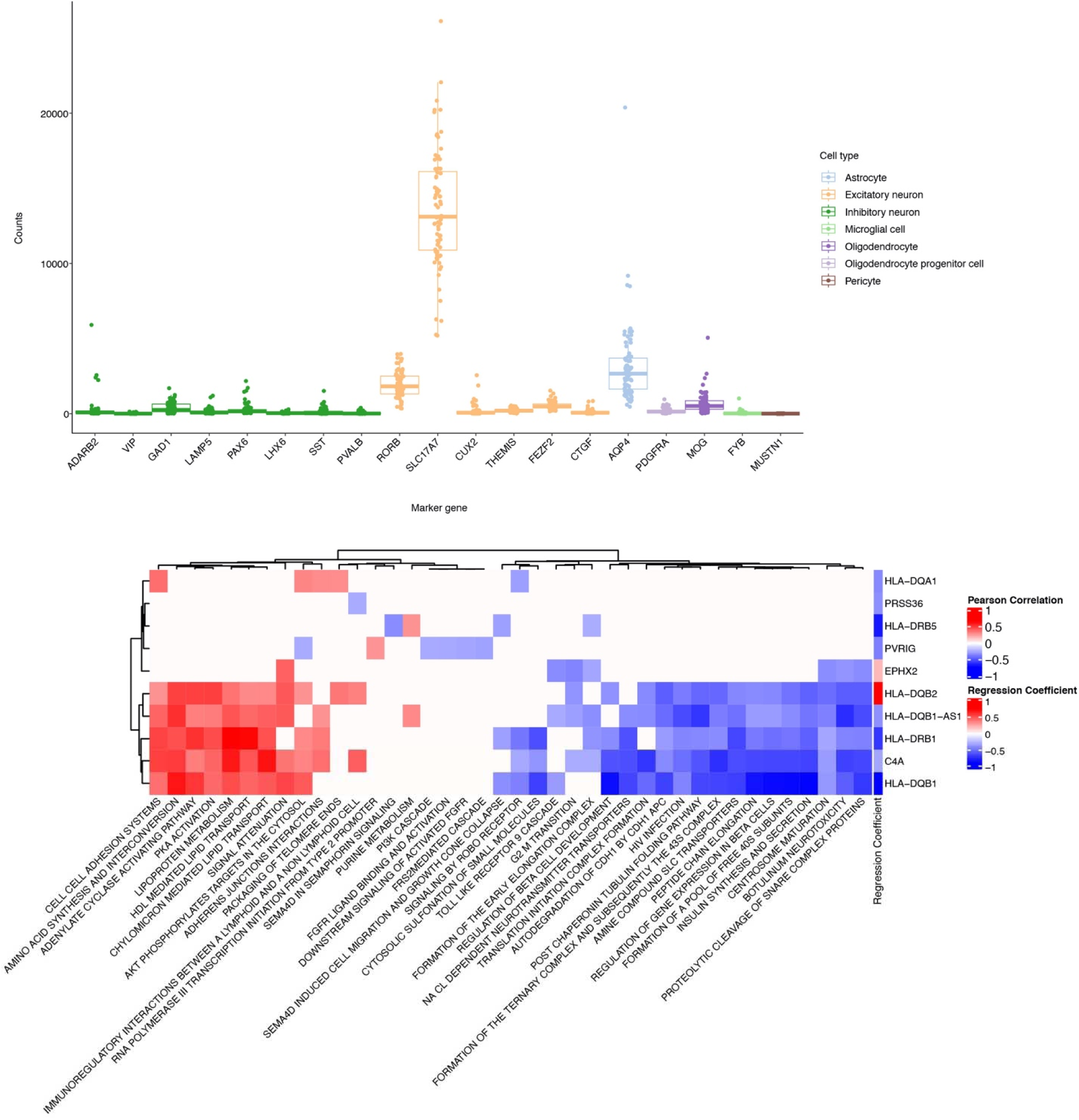
a) Boxplots depict the normalized counts of cell type marker genes in the brain in the temporal cortex laser capture microdissected neurons. Boxes are colored by the general cell type represented by the marker gene. Each point represents one bulk RNA-seq sample. b) Heatmap showing significant Pearson correlations between the RNA-seq expression of temporal cortex pyramidal neuron eGenes and Gene Set Variation Analysis signatures for REACTOME pathways. The gene names on the rows are annotated for the regression coefficient representing the association between gene expression and the presence of the associated Alzheimer’s disease eQTL. The legend for these regression coefficients is labeled as “Regression Coefficient”. Columns are clustered with hierarchical clustering.

**Extended Data Fig. 4, related to Fig. 3.**
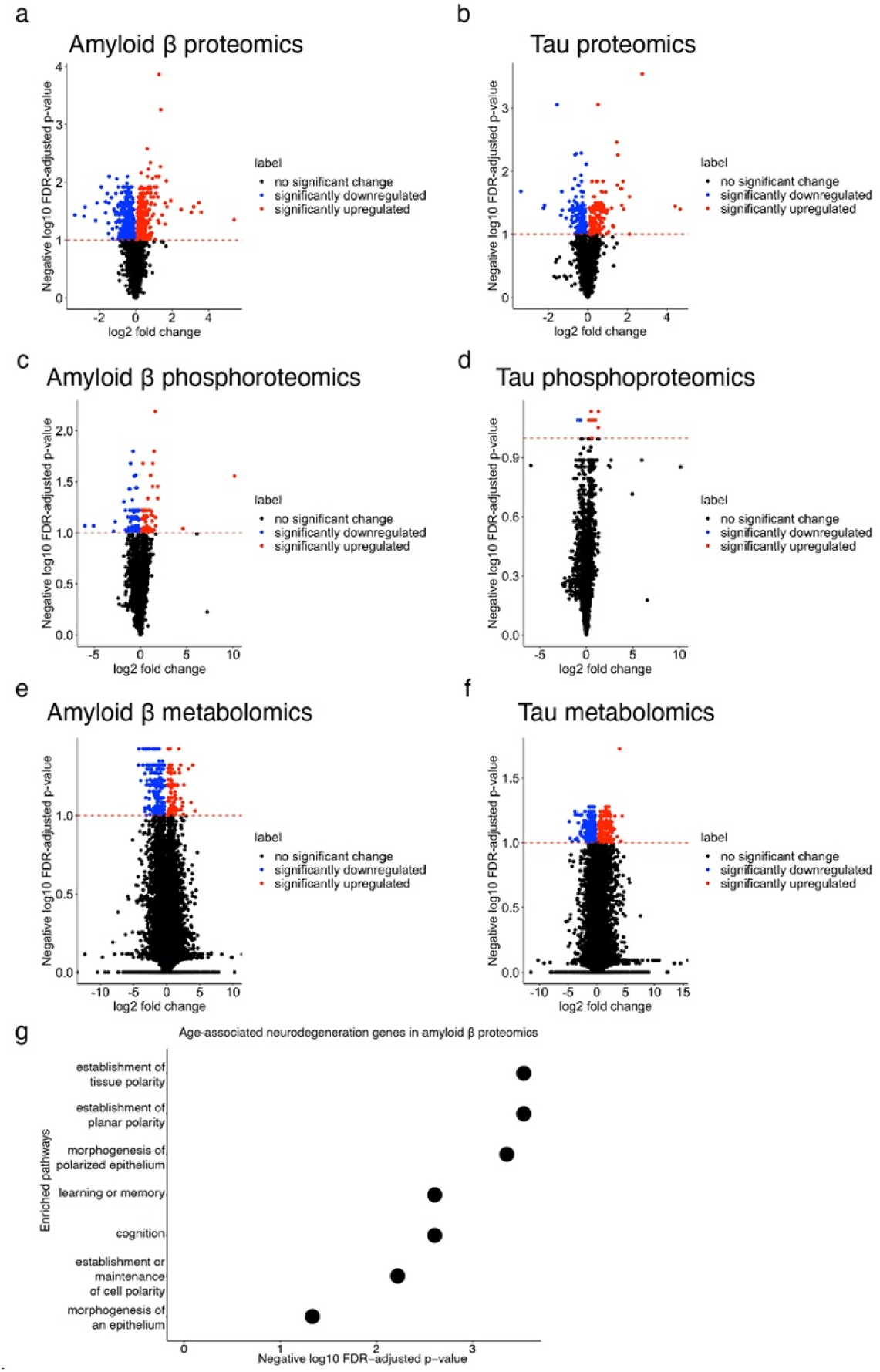
Volcano plots depicting negative log_10_ FDR-adjusted p-values and log_2_ fold changes between case and control in a) proteomics from Aβ_1-42_ transgenic flies (amyloid β), b) proteomics from tau^R406W^ transgenic flies, c) phosphoproteomics from Aβ_1-42_ transgenic flies, d) phosphoproteomics from tau^R406W^ transgenic flies, e) metabolomics from Aβ_1-42_ transgenic flies, and f) metabolomics from tau^R406W^ transgenic flies Blue dots indicate significantly downregulated omics and red dots indicate significantly upregulated omics. The horizontal red dashed line indicates the FDR cut-off at 0.1. g) Dot plot indicating GO terms overrepresented in neurodegeneration screen hits that are differentially abundant in proteomics from Aβ_1-42_ transgenic flies.

**Extended Data Fig. 5, related to Fig. 5:**
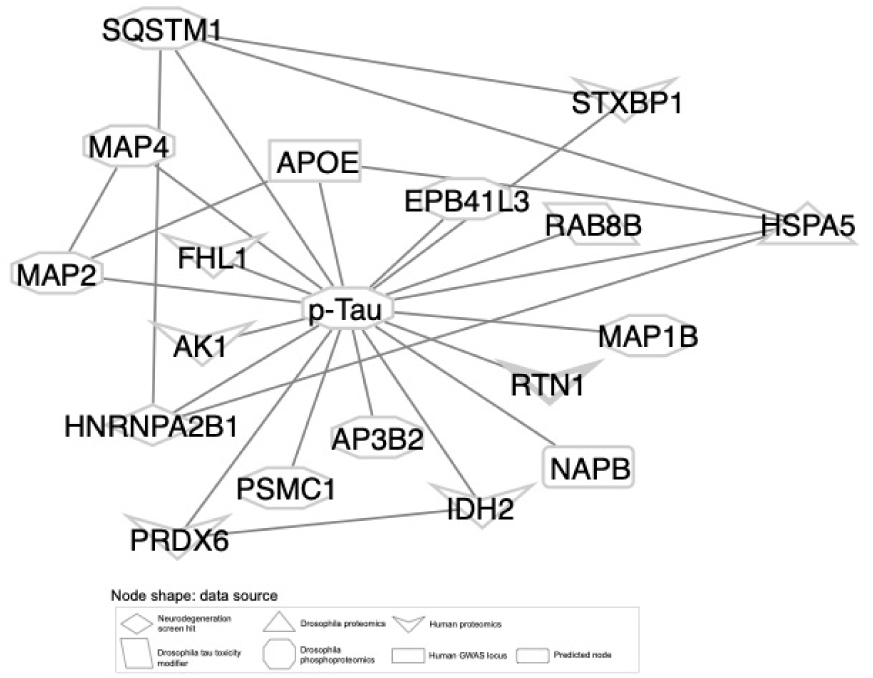
pTau and its first neighbors in the network solution. Nodes are shaped by the data source from which they were contributed.

**Extended Data Fig. 6, related to Figs. 5 and 6:**
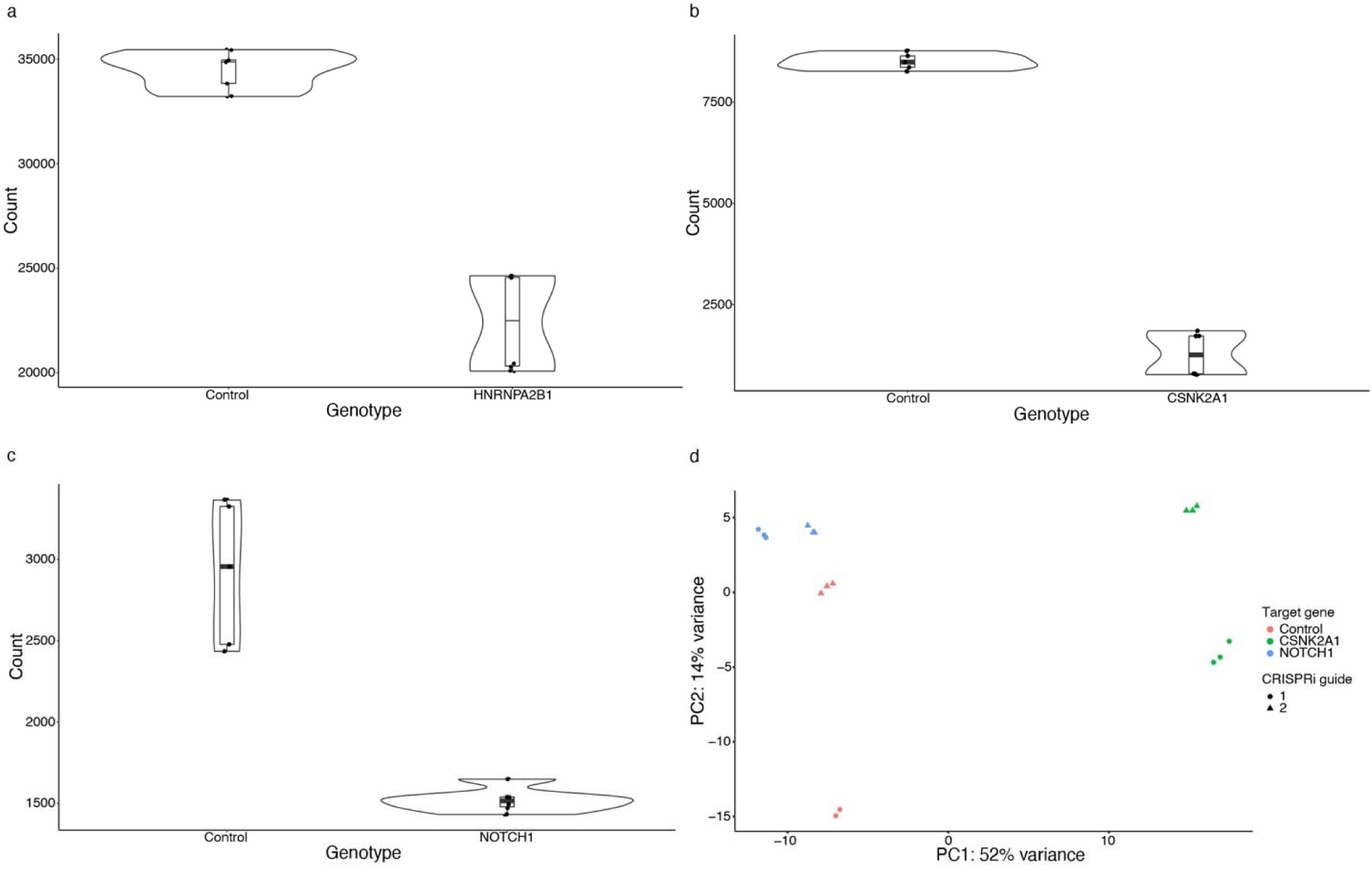
Knockdown efficiency and principal component analysis of the NGN2 RNA-seq data. Violin plots depict library-corrected RNA-seq counts in NGN2 neuronal progenitor cells for controls and a) *HNRNPA2B1*, b) *CSNK2A1* or c) *NOTCH1* knockdown. d) Principal Component Analysis plot of individual control, *NOTCH1* and *CSNK2A1* RNA-seq replicates from expression data. Colors indicate the knockdown for each replicate and the shape indicates whether the knockdown was performed with the first or second guide RNA. For control, we used non-targeting guide RNAs.

**Extended Data Fig. 7, related to Fig. 6:**
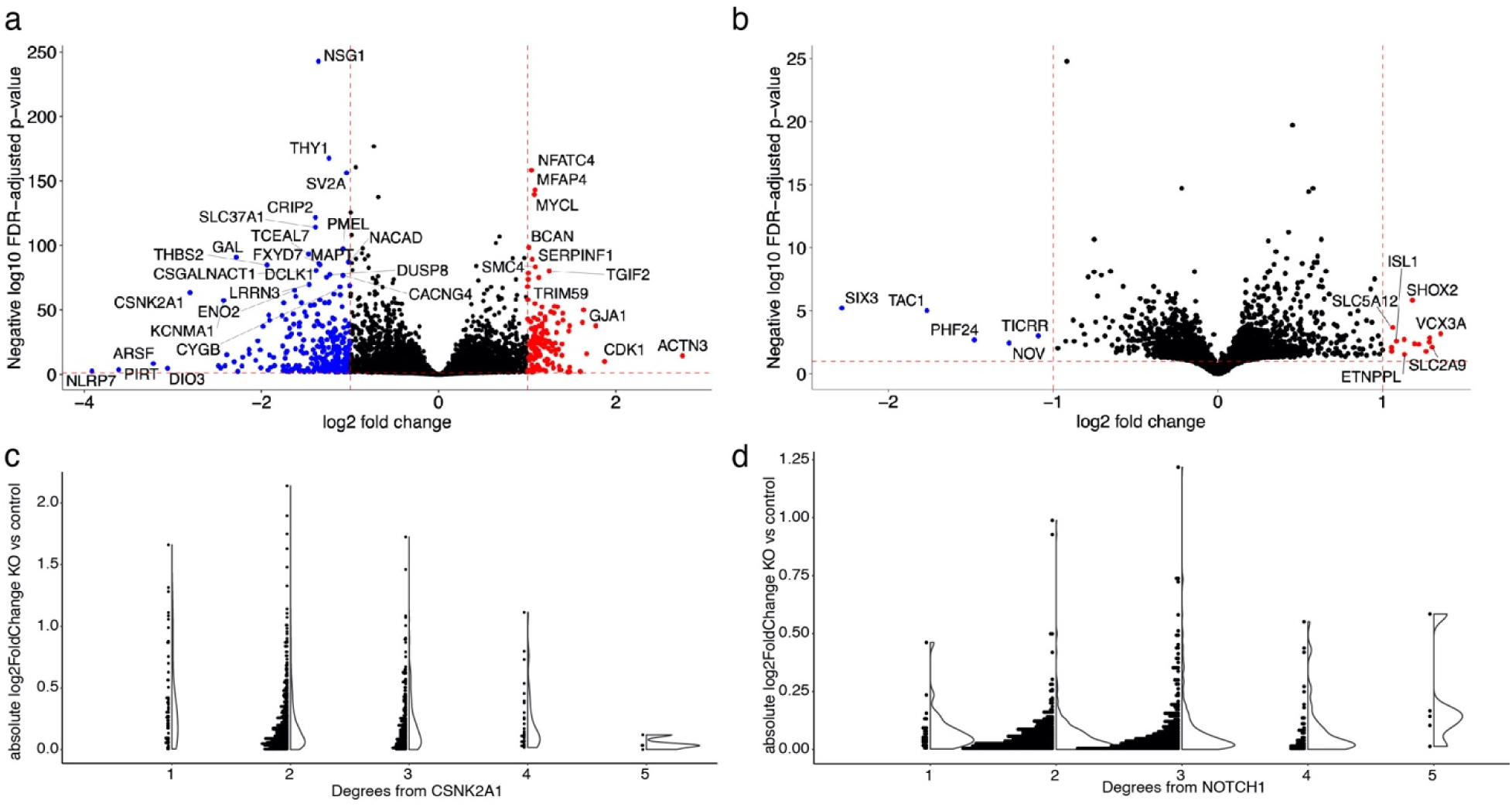
RNA-seq analysis after knockdown of *NOTCH1* and *CSNK2A1*. Volcano plot depicts differential expression analysis by DeSeq2 of bulk RNA-seq after a) CSNK2A1 CRISPRi knockdown and b) NOTCH1 CRISPRi knockdown in NGN2 iPSC-derived neural progenitor cells. Each dot represents a single gene. The horizontal dashed line indicates the negative log_10_ Benjamini-Hochberg FDR-adjusted p-value cut-off of 0.1 and the vertical dashed lines indicate the log_2_ fold change cut-offs of 1 and -1. Red dots indicate significantly upregulated genes (log_2_ fold change greater than 1) and blue dots indicate significantly downregulated genes (log_2_ fold change less than -1). c) Dot and volcano plots show the absolute value of the log_2_ fold change of nodes in the network after *CSNK2A1* knockout compared to controls relative to the degree of separation from *CSNK2A1*. d) Dot and volcano plots show the absolute value of the log_2_ fold change of nodes in the network after *NOTCH1* knockout compared to controls relative to the degree of separation from *NOTCH1*.

